# When and why modular representations emerge

**DOI:** 10.1101/2024.09.30.615925

**Authors:** W. Jeffrey Johnston, Stefano Fusi

**Affiliations:** Center for Theoretical Neuroscience; Mortimer B. Zuckerman Mind, Brain, and Behavior Institute; Kavli Institute for Brain Science Columbia University, New York, NY, USA

## Abstract

Experimental and theoretical work has argued both for and against the existence of specialized sub-populations of neurons (modules) within single brain regions. By studying artificial neural networks, we show that this local modularity emerges to support complex (for instance, context-dependent) behavior only when the input to the network is low-dimensional. No anatomical constraints are required. We also show when modular specialization emerges implicitly at the population level, where orthogonal subspaces are viewed as distinct modules. Modularity yields abstract representations, allows for rapid learning and generalization on novel output domains as well as related tasks. Non-modular representations facilitate the rapid learning of unrelated tasks. Our findings reconcile conflicting experimental results and make predictions for future experiments.

## 1 Introduction

The organizational principles of neural activity within single brain regions are still largely unknown. Some work provides evidence for the existence of specialized subpopulations of neurons that represent specific variables[1, 2] or that are active in specific contexts[3, 4]. Other work[5, 6] argues that these specialized neurons are the exception, rather than the rule, where most neural populations can be described by a continuous distribution of tuning across different task conditions. Both forms of single neuron selectivity can produce neural population representations with the computational advantages of both low- and high-dimensional representations, consistent with recent experimental work[7–13]. While previous theoretical work has shown that specialized subpopulations of neurons emerge naturally due to constraints on the connectivity and physical arrangement of neurons on the cortical sheet[14–16] – thus providing an explanation for large-scale, region-level organization in the brain[17–20] – we lack an explanation for why specialized subpopulations of neurons emerge within a single brain region in some cases[1–4] but not others[6–8, 10, 11, 13, 21, 22]. To investigate this question, we focus on two components of a learning system: The task at hand, which is often complex and nonlinear with respect to experimenter-defined latent variables[22–26], and the dimensionality – or representational geometry – of the neural representations at the start of training, which is often unconstrained by as well as unobserved in neuroscience experiments.

For complex tasks, specialized subpopulations of neurons can group related task conditions together and simplify computation. Some laboratory tasks have natural condition groupings, such as context-dependent behavior. These tasks often involve applying a simple rule within each individual context, but gain complexity because different rules must be applied across different contexts. For instance, when shopping for a banana to eat right away, the subject will search for yellow bananas – a simple, color-based discrimination. However, if the subject is shopping for a banana to eat later in the week, they will search for a green banana. Thus, context defines how the same visual stimuli are acted upon. A neural solution to this kind of task could consist of one population of neurons that is active only when searching for a banana to eat now and a distinct population of neurons that is active only when searching for a banana to eat later in the week. This population level structure effectively decouples the two behaviors and allows a downstream brain region to solve the full task by learning to decode the ripeness of the fruit from whichever population of neurons is active at a given time. We refer to this as an *explicitly modular* representation – and there is experimental evidence that this kind of structure exists in some cases[3, 4]. This kind of decomposition can also occur at the population level, without requiring specialized subpopulations of neurons. Instead, the neural population activity can be confined to orthogonal subspaces during each of the two contexts. We refer to this as an *implicitly modular* representation – there is also experimental evidence that this kind of structure exists in other cases[22, 26–29]). Implicit modularity has the same computational advantages as explicit modularity (i.e., a downstream neuron can easily read out one module and ignore the others).

However, we show that the computational benefit of modularity described above only holds when a very specific – but common – assumption about the input to the neural system is made: That the idealized variables designed by the experimenter are provided directly as input to the model[24–26, 30]. From experimental data, we know that this assumption is typically violated[7–13], as distinct variables are almost always nonlinearly mixed with each other (e.g., neurons that respond only to specific combinations of certain variables, like shape and color)[12, 21, 31]. This mixing yields a much higher dimensional input than typically assumed by modeling studies. High-dimensional input representations allow even complex tasks to be solved directly by a simple readout (such as a single neuron or a linear decoder) without requiring either explicit or implicit modularity[21, 31]. Thus, we hypothesize that the dimensionality of the input representation will shape the kind of representation that is learned to solve a particular task.

Here, we confirm this hypothesis for a wide variety of tasks. In particular, we show that the strong explicit modularity observed in previous work[24–26, 32] emerges in neural networks only for low-dimensional input representations, while high-dimensional unstructured input representations yield learned representations that are either unstructured or implicitly modular depending on the number of task output dimensions. We show that the transition between these two regimes depends on the number of stimulus features. Next, we focus on context-dependent behavior and characterize how the learning of novel contexts depends on input structure and the number of outputs. For novel contexts that are related to previously learned contexts, we find that modular representations speed learning; while non-modular representations speed the learning of novel but unrelated contexts. Finally, we generalize our framework to a broad class of tasks, and show which lead to modularity as well as verify that input structure determines whether modularity emerges. Our theory provides a potential explanation for diverse experimental findings related to modularity. Further, they provide predictions both for the representational geometry that will be observed in novel experimental conditions as well as for the ability of animals to generalize and learn in novel contexts. We close by discussing these predictions in more detail.

## 2 Results

We want to understand how representations of stimuli described by multiple features are shaped by both the input geometry and task demands. While we study this in artificial neural networks, we will make predictions for real neural data and provide a theoretical understanding of the behavioral and computational constraints that shape neural population representations in the real brain.

To do this, we focus on what we term as complex tasks, which we define as tasks that are not linearly separable with respect to the variables on which they depend. As an example task, consider shopping for a banana. The decision to buy a banana or not depends on both the color of the banana and whether the intention is to eat it now or later (fig. 1a). The intention changes the behavioral meaning of the color: when the intention is to eat the banana now, yellow bananas are desirable; when the intention is to eat the banana in a few days, green bananas are desirable. This task is an example of a broader class of tasks that cannot be solved directly by a simple linear decoder (fig. 1a, right), which must find a linear boundary in the space described by the two latent variables that accurately determines the correct action (i.e., select or avoid). This task in particular is an example of the canonical exclusive-or relation between two variables, which is a well-known non-linearly separable problem with applications to neuroscience[31].

**Figure 1:**
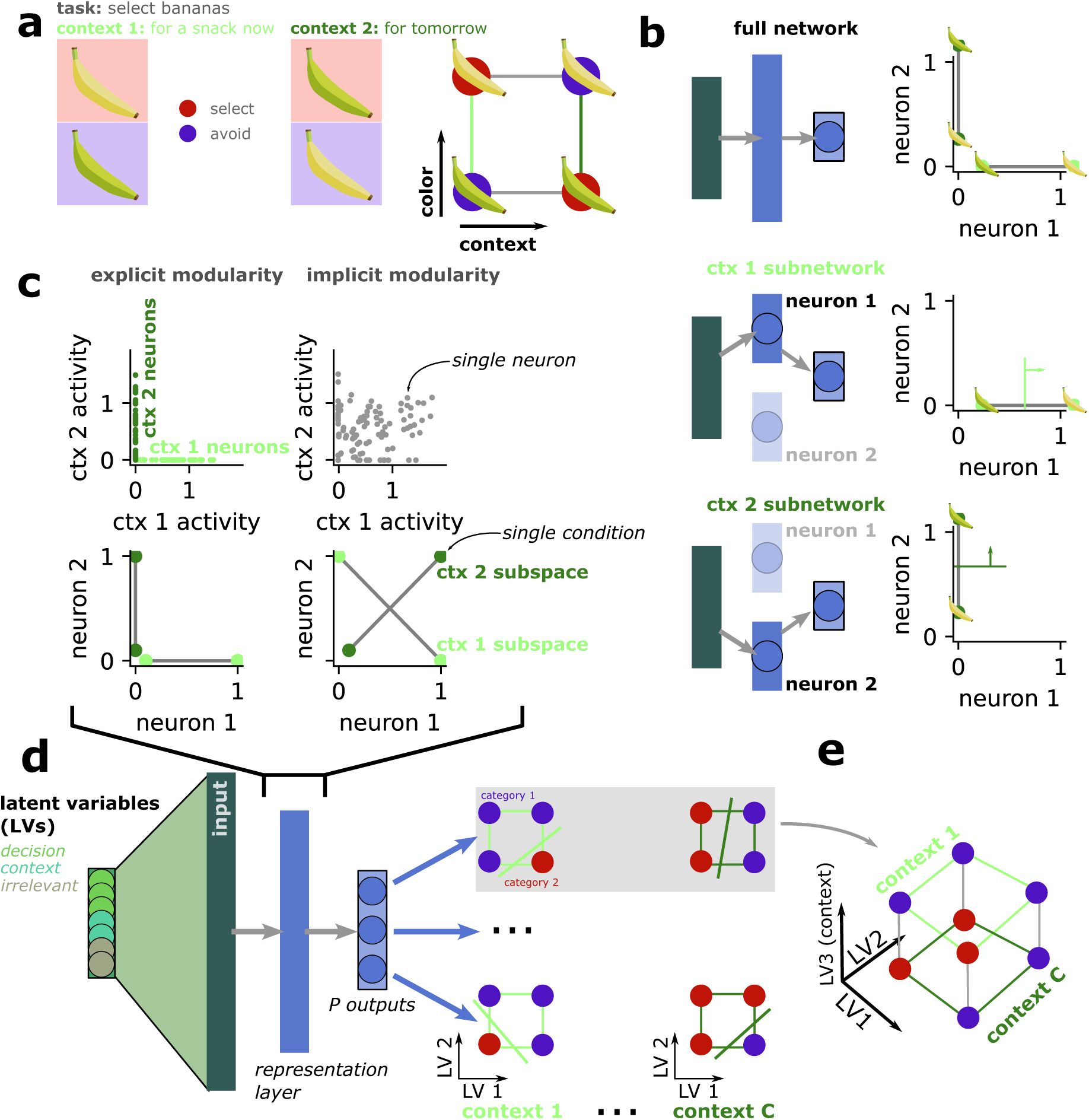
Contextual behavior and modularity in artificial neural networks. **a** An example contextual task on banana stimuli (left) as well as a low-dimensional representation of the two variables (right). **b** A modular solution to the contextual task in a neural network. (top) The neural network has two neurons, one dedicated to each of the two contexts. (middle) In the first context, only the first neuron is active and the output layer (one neuron in this simple case) can decode the appropriate action based on the activity of that neuron while ignoring the other neuron. (bottom) In the second context, only the second neuron is active and the output layer can decode the appropriate action based on the activity of that neuron while ignoring the other neuron. **c** Schematic of explicit and implicit modularity. For explicit modularity, single neurons are specialized for one of the two contexts (top left). At the population level, this means there are two subspaces, one for each context, aligned with different neurons (bottom left). For implicit modularity, there is no specialization at the level of single units (top right), but at the population level there are still specialized and orthogonal subspaces for each context (bottom right). **d** Schematic of the feedforward neural network used here. The network receives representations of decision, context, and irrelevant variables through an input model (described below). The network has a single hidden layer (representation layer) and an output layer, where there are *P* output units. In each context, every output unit must learn a linear classification of the decision variables. **e** Combined across multiple contexts, the output required from each output unit is not linearly separable given a low-dimensional input representation.

This task is also an example of the first class of tasks that we will focus on in this work: A contextual task. In this case, the intention can be viewed as a context variable. This contextual structure has a natural connection to explicit modularity, in which distinct subpopulations of neurons are active for each of the different behavioral contexts. In particular, while the full contextual task cannot be solved by a single linear decoder (fig. 1a, right), a nonlinear feedforward network can solve the full task by decomposing it into linearly separable sub-tasks according to context (fig. 1b). Each sub-task is defined by a subset of conditions and can be solved by an independent sub-network of the full network (fig. 1b, middle, bottom) without interference between the different contexts. Indeed, each sub-network can be trained independently to solve a specific sub-task as the synapses of the inactive neurons are ignored. So the original non-linearly separable problem can be decomposed into simpler, linearly separable sub-tasks. In our banana example, each sub-task contains only two conditions, which are always linearly separable. The sub-network structure can be either explicit (fig. 1c, left) or implicit (fig. 1c, right).

We will study when and how explicit and implicit modularity (fig. 1c) emerge in artificial neural networks trained to perform context-dependent tasks like this example task. The complex task network receives decision (e.g., color) and context variables (e.g., whether buying a banana for now or for later) alongside irrelevant variables (e.g., where the bananas are in the store; fig. 1d, left). The network is trained to solve a contextual task that has one or more outputs (fig. 1e). Each output is linearly separable within a single context (fig. 1e, “context 1”), but will – on average – not be linearly separable across multiple contexts (fig. 1e). All outputs share the same contextual structure, but differ in the linear classification boundary that must be learned within each context (fig. 1d, top right relative to bottom right). To investigate when and how these different representational strategies emerge, we will vary the geometry of the input representations provided to the neural network as well as the number of outputs that the network must produce. Later, we will move away from this strict contextual structure to show how our findings generalize to arbitrary task structures.

### 2.1 The representational geometry of the input

Our input always consists of several decision-related variables (*D* = 3, several context variables (*C* = 2), and several irrelevant variables (*I* = 4; see *Input model* in *Methods*). We hypothesize that the geometry of the input representations provided to the complex task network is one of the primary factors that shapes the learned representations. To explore this idea, we develop an input model that interpolates between a low-dimensional, disentangled input representation with the same embedding dimensionality as the number of latent variables (*D* + *C* + *I*, fig. 2a) and a high-dimensional, conjunctively mixed, unstructured input representation with an embedding dimensionality that scales exponentially with the number of latent variables (2*^D^*^+*C*+*I*^, fig. 2b). To smoothly interpolate between these two extreme cases (fig. 2c), the full input is constructed from a weighted sum of the two extremes. As the weight parameter (fig. 2d, *s*) varies from 1 to 0, the input structure transitions from fully disentangled to fully unstructured and increases in dimensionality (fig. 1c, and see *Dimensionality of unstructured solutions* in *Methods*). For the low-dimensional, fully disentangled input, each input variable is provided along a separate dimension of the input space. In the brain, the linear tuning for facial features in the inferotemporal cortex is consistent with a disentangled representation[33, 34]. In the unstructured input, all input variables are nonlinearly mixed together, such that each unique stimulus (defined as a particular combination of decision, context, and irrelevant variables) is represented along an orthogonal dimension in the input population space. For instance, instead of having units that respond to color and size independently – as in fig. 2a – there are units that respond only to particular combinations of color, size, and other variables – as in fig. 2b. In the brain, many areas of prefrontal cortex have been shown to have highly nonlinear mixed representations[21, 31]. In between these two extremes are representations with both disentangled and nonlinear representations superimposed on each other (fig. 2d). Previous theoretical and empirical work has shown that such intermediate representations can preserve the computational benefits of both disentangled and high-dimensional representations[8, 10, 13], as well as shown that many brain regions in sensory and frontal cortex exist in this middle ground [7–10, 13]. We will show that changes to this input structure will qualitatively change the representations learned by networks performing complex tasks.

**Figure 2:**
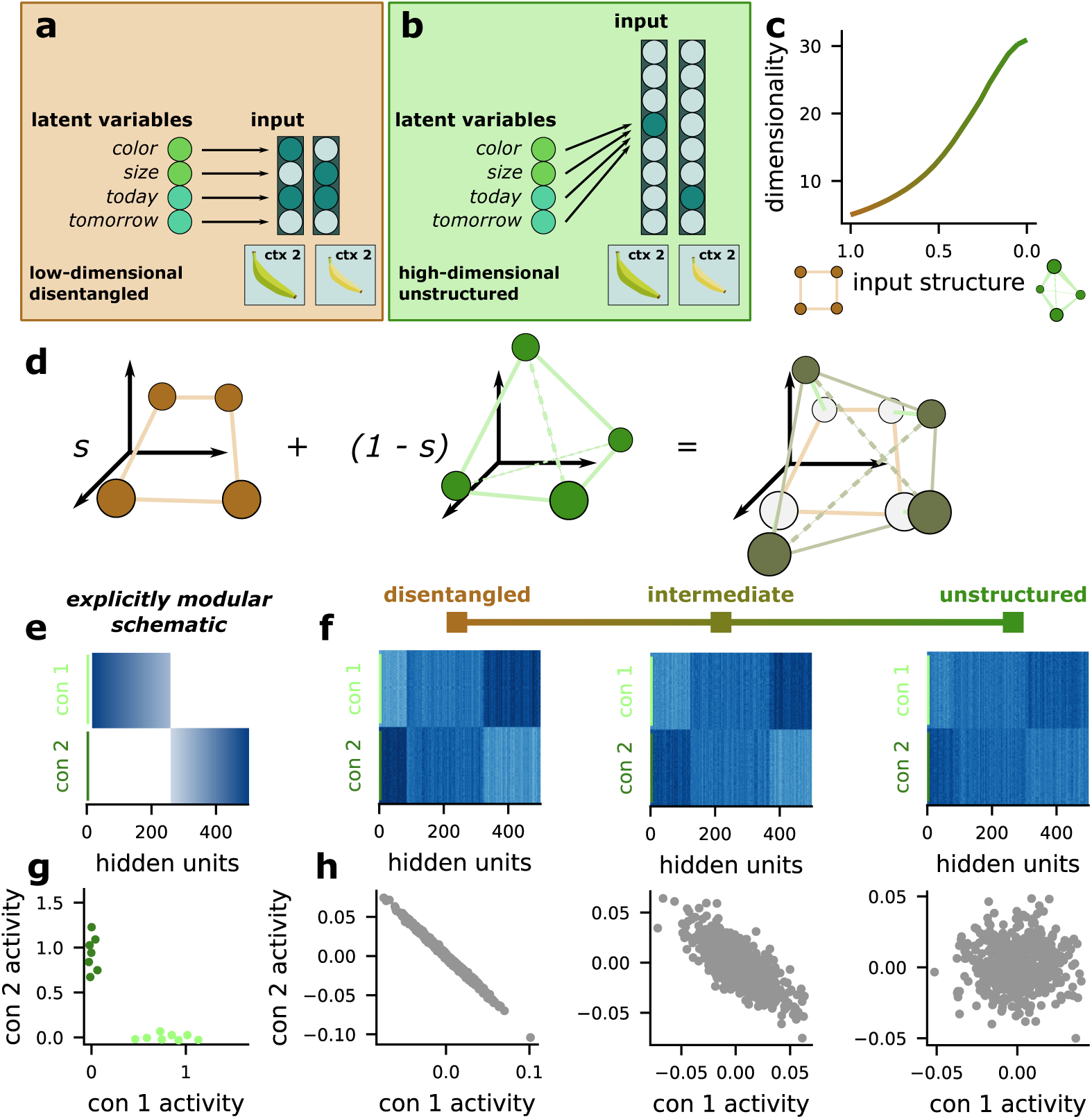
Varying the representational geometry produced by the input model. **a** The low-dimensional, disentangled input model has the different latent variables confined to different dimensions in input space. **b** The high-dimensional, unstructured input model has nonlinear, conjunctive mixed selectivity for all latent variables, such that the input associated with each stimulus is equivalent to a one-hot vector. **c** The extreme cases in **a** and **b** describe a spectrum of input structure, where the embedding dimensionality of the input varies smoothly from low to high. **d** We produce input models between these two extremes by superimposing input representations with these two different geometries onto each other, where the two components are given different weights depending on their position on the spectrum. **e** A schematic showing the expected pattern of responses for a representation with explicit modularity when the y-axis is sorted by the context from which a stimulus is sampled and the x-axis is sorted by the difference in average activity between the two contexts. **f** The same analysis in **e** applied to input models from different points on the spectrum. **g** A schematic showing the expected pattern of single unit average activity in each context for a representation with explicit modularity. **h** The same analysis in **h** applied to input models from different points on the spectrum.

### 2.2 Measures of modularity

We develop two methods for visualizing these representations that will reveal modular structure if it exists. We introduce these visualizations now, since they are used to analyze the representations learned by the complex task network later. First, we sample stimuli from each of the two contexts (fig. 2e, y-axis) and sort the units in the input representation by the difference in average activity between the two contexts (fig. 2e, x-axis). A modular representation would have a clear block structure under this analysis (fig. 2e). This sorting procedure reveals a weak block structure for all of the different input geometries that we consider (fig. 2f), which could reflect pre-existing modular structure within our input model. To investigate this structure further, we compute the average activity in each context for every unit in our input model. Then, we cluster the units in this mean activity space, and color them according to their cluster membership (fig. 2g). We choose the number of clusters according to the Bayes information criterion (see *Clustering analysis* in *Methods* for details). In a model with strong modular structure related to context, the units should cluster into two groups along the axes of this space, where one subset of units have non-zero average firing rates in the first context (but not the second) while a distinct subset of units have non-zero average firing rates in the second context (but not the first; fig. 2g). In the input model, we find the points are best explained by a single cluster for each input geometry (fig. 2h), indicating that there is no modular structure in any of our input models, despite the apparent structure present in the earlier visualization (fig. 2f). The disentangled input models show a negative correlation in average activity across the two contexts. This is because the contexts themselves are negatively correlated: for each stimulus, one context variable will be one, and the other will be zero.

### 2.3 The complex task network

The complex task network (fig. 3a) receives the input representations produced by the input model (fig. 3a) and is trained to produce *P* labels of each stimulus along different output dimensions, where the labels are determined by *P* randomly chosen binary classification tasks (fig. 3c). We assume that within each context, all output labels are linearly separable with respect to the original latent variables (and they are also linearly separable with respect to our input representations). However, in most cases, the full task (combined across both contexts) will not be linearly separable (as discussed above, fig. 1c, bottom left). The readout layer of the network is effectively a set of linear classifiers. So, we know that, if the network successfully produces all *P* output labels, then those *P* classifications must be made linearly separable in the hidden layer. Both forms of modular representation introduced above satisfy this constraint.

**Figure 3:**
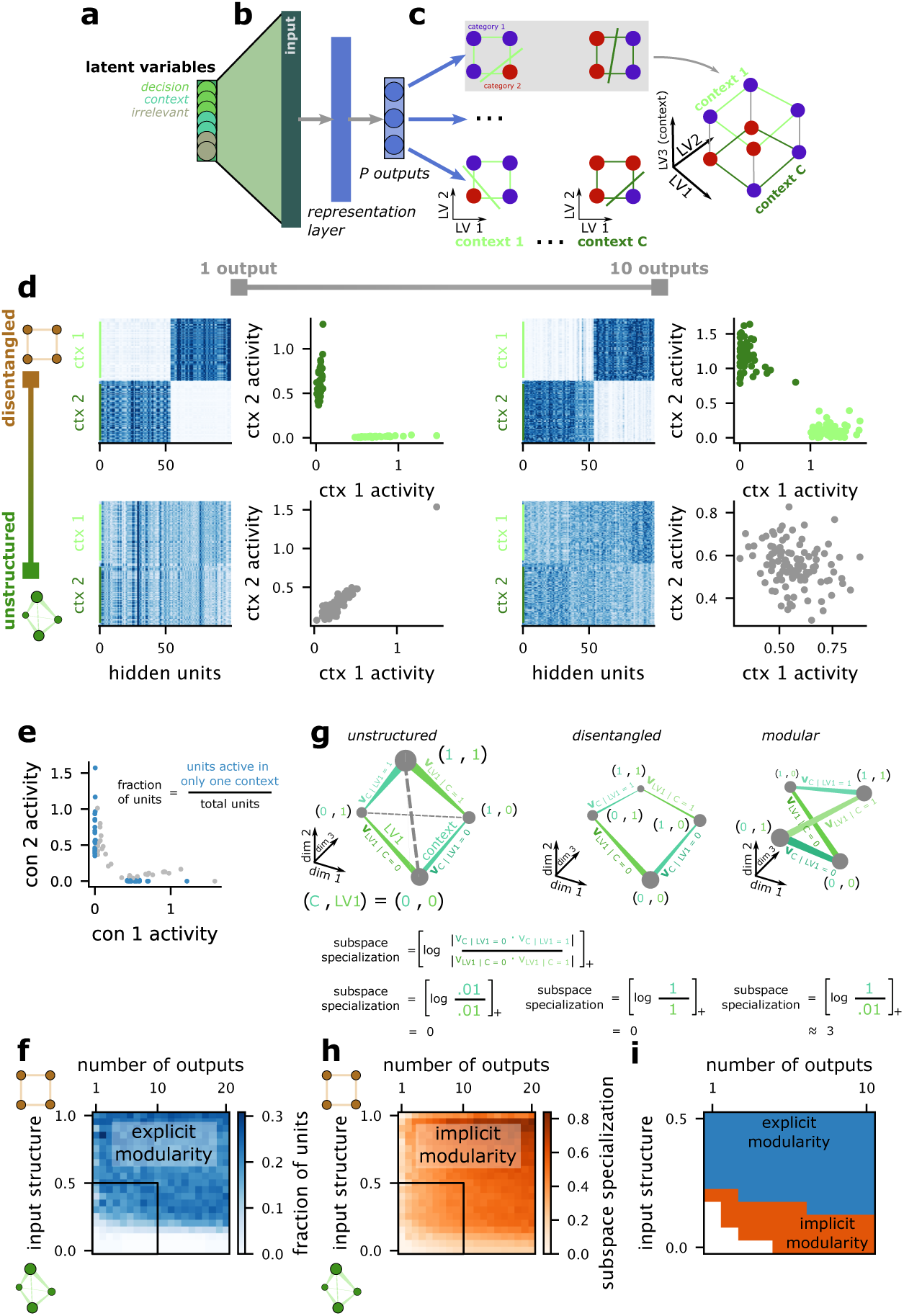
Modular representations emerge when the input is disentangled or the network is multi-tasking within each context. **a** The input model, already described in Fig. 2. **b** The contextual multi-tasking model is a single hidden-layer feedforward network trained to perform contextual tasks with *P* output dimensions across *C* = 2 contexts. **c** Each task is a linear classification task within each context (left). The linear boundary is randomly selected for each context, giving rise to a nonlinear classification task across multiple contexts (right). **d** Hidden layer representational structure across several example models. The hidden unit activations in both contexts, sorted by cluster membership (left). The average activity of each unit in the two contexts, colored according to cluster membership (right). The models in the left column are trained to perform *P* = 1 task in each context, while the models in the right column perform contextual tasks with *P* = 10 output dimensions on the same 3 latent variables in each context. The models in the top row are given disentangled inputs; the models in the bottom row are given high-dimensional, unstructured inputs. **e** The fraction of units active in only a single context given different choices for the number of output dimensions *P* and the input geometry. **f** The fraction of units specialized for a single context (defined as in **e**) for the full parameter range. Specialized units are indicative of explicit modularity (top). **g** The subspace specialization metric applied to three different cases (see *Subspace specialization* in *Methods* for more details). The measure is zero for both unstructured (left) and disentangled (center) representations of the variables. It is positive for a modular representation (right). **h** The same as **f** except showing the subspace specialization metric, a measure of implicit modularity. **i** The reduced parameter range indicated by the black box in **f** and **h**. Blue shows where the fraction of specialized units is *> .*05, while orange shows where subspace specialization is *> .*05 out of the remaining parameter choices. White has unstructured representations.

We begin by visualizing the activity in four trained networks that vary in both the input geometry and in the number of output dimensions that the complex task network is trained to use. First, we train a network to perform a contextual task with a single output *P* = 1 across *C* = 2 contexts, where the same *D* = 3 latent variables are relevant in both contexts. We train this network with fully disentangled input representations (fig. 3d, top left). This network learns explicitly modular representations, in which some units in the hidden layer are active in one of the two contexts but not the other. The units in the hidden layer cluster into two broad groups: one subset of units are active in the first context but relatively silent in the second (fig. 3d, dark green); a distinct subset of units is active in the second context but relatively silent in the first (fig. 3d, light green and gray). We quantify modular structure by computing the fraction of units in the hidden layer that are active in one context but nearly silent in the other (fig. 3e, and see *Contextual fraction* in *Methods* for details). This pattern of results holds when we increase the number of output dimensions for the contextual task (*P* = 10, fig. 3d, top right). The activity in these networks resembles neural activity recorded from posterior parietal cortex in a mouse performing a contextual discrimination task[3].

Next, we train a network to perform a contextual task with one (*P* = 1, fig. 3d, bottom left) and ten (*P* = 10, fig. 3d, bottom right) output dimensions on fully unstructured input representations. This eliminates the explicitly modular structure from before: Neither network develops significant clustering in average activity space. Next, we apply our metric for explicit modularity to networks trained on a full range of input geometries and numbers of tasks (fig. 3f). This analysis reveals that functionally specialized sub-populations of units (i.e., explicit modularity) emerge as a function of the input geometry and this emergence has only a weak dependence on the number of outputs. A critical value of nonlinear mixing in the input geometry marks a transition between modular and non-modular solutions to the contextual tasks. We show how this critical value depends on the features of the input, and link the emergence of modularity to a competition between learning from the disentangled and unstructured components of the full input representation (see *Explicit modularity emerges when the disentangled component of the input provides faster learning* in *Supplement* and fig. S3).

However, there could be additional structure in the learned representations – that is, this fraction of contextual units only quantifies explicit modularity, but does not quantify implicitly modular structure. To quantify this, we develop a metric that we refer to as subspace specialization (fig. 3g). This metric depends on the overlap between subspaces defined by different splits of the condition space into two halves, which we measure using the alignment index[35]. First, we compute the overlap between the subspace containing all the conditions from one context and the subspace containing all the conditions from the other context. Then, we compare that context overlap to the overlaps we obtain by treating the decision variables as if they were the context variable and repeating the analysis. That is, for a given decision variable, we split all stimulus conditions into two sets: for one set, the decision variable has one value; for the other set, it has its other value.

Then, we take the overlap between the subspaces containing these two distinct sets of points. This gives us a set of decision variable overlaps to compare to the context variable overlap. If both the decision variable and context overlaps are low, subspace specialization is low (fig. 3g, left); the representation is similar to our unstructured input. If both the decision variable and context overlaps are high, subspace specialization is also low (fig. 3g, middle); the representation is similar to our disentangled, low-dimensional input. However, if the overlap between the two context-specific subspaces is low, but the decision variable overlaps are high, then subspace specialization is high (fig. 3g, right). In particular, subspace specialization will be positive for representations that split into orthogonal subspaces when conditioned on a contextual variable but not on other variables (fig. 3g, right, and see *Subspace specialization* in *Methods* for a full description of the metric).

Using subspace specialization, we quantify the emergence of implicitly modular representations as a function of input geometry and the number of output dimensions for the contextual task (fig. 3h, right). As expected, whenever a representation is explicitly modular, it also has positive subspace specialization. However, implicit modularity also emerges for high-dimensional input geometries once the network is trained to use a sufficient number of output dimensions. Thus, the only regime without explicit or implicit modularity is for complex task networks performing a contextual task with few output dimensions and high-dimensional inputs. To illustrate these transitions, we focus on a reduced parameter range where networks have relatively unstructured inputs and the contextual tasks have relatively few output dimensions (fig. 3f, h, black box). Then, we binarize both measures used above, and show the transitions between the three representational regimes (fig. 3i).

This framework can help to explain the heterogeneous representations observed for contextual behavior in the brain. It reveals that the representational geometry learned by the network depends strongly on the geometry of the input representations – or, alternatively, on the representational geometry that exists prior to the start of training on the new task – as well as on the dimensionality of the behavior.

### 2.4 The learned modular structure facilitates rapid learning

We show that the representations learned by the complex task networks within each context are abstract (see *The model develops an abstract and high-dimensional representational geometry* in *Supplement* and fig. S4), consistent with previous work[36]. These abstract representations have been widely observed in the brain[8–11, 37] and enable the learning of novel output associations from only a handful of stimulus examples[10, 36]. We show that the level of abstraction in the learned representation depends primarily on the number of outputs learned by the network: more outputs give rise to more abstract representations (fig. S4), consistent with prior work[36]. In contrast, we find that zero-shot generalization to novel stimuli – a distinct, but related form of generalization – depends primarily on the input structure (fig. S5). In this case, disentangled input representations allow reliable zero-shot generalization, while unstructured input representations – even with many output dimensions – give rise to chance-level zero-shot generalization performance. Thus, complex task network trained with multiple outputs and disentangled input representations will exhibit both reliable zero-shot generalization as well as rapid learning of new outputs.

Next, we study how the input geometry shapes the learning of a novel task in two previously learned contexts (fig. 4a). In this case, we train each network to perform contextual tasks with *P* output dimensions in *C* = 2 contexts. Then, after training on the original task, we introduce a *P* + 1th output to the task – which depends on the same set of variables – and train the network to perform this supplemented task. Then, we measure how task performance changes across training on this new output dimension (fig. 4b, top). We show that the learning speed for a new output depends on both the input geometry and the number of output dimensions in the original task (fig. 4b, bottom). That is, both lower-dimensional input geometries and a larger number of previously learned output dimensions increase the speed with which the new output is learned. This is consistent with previous work showing that learning from lower-dimensional representations is typically faster than learning from higher-dimensional representations[36, 38], so long as the low-dimensional representations allow for the classification to be learned.

**Figure 4:**
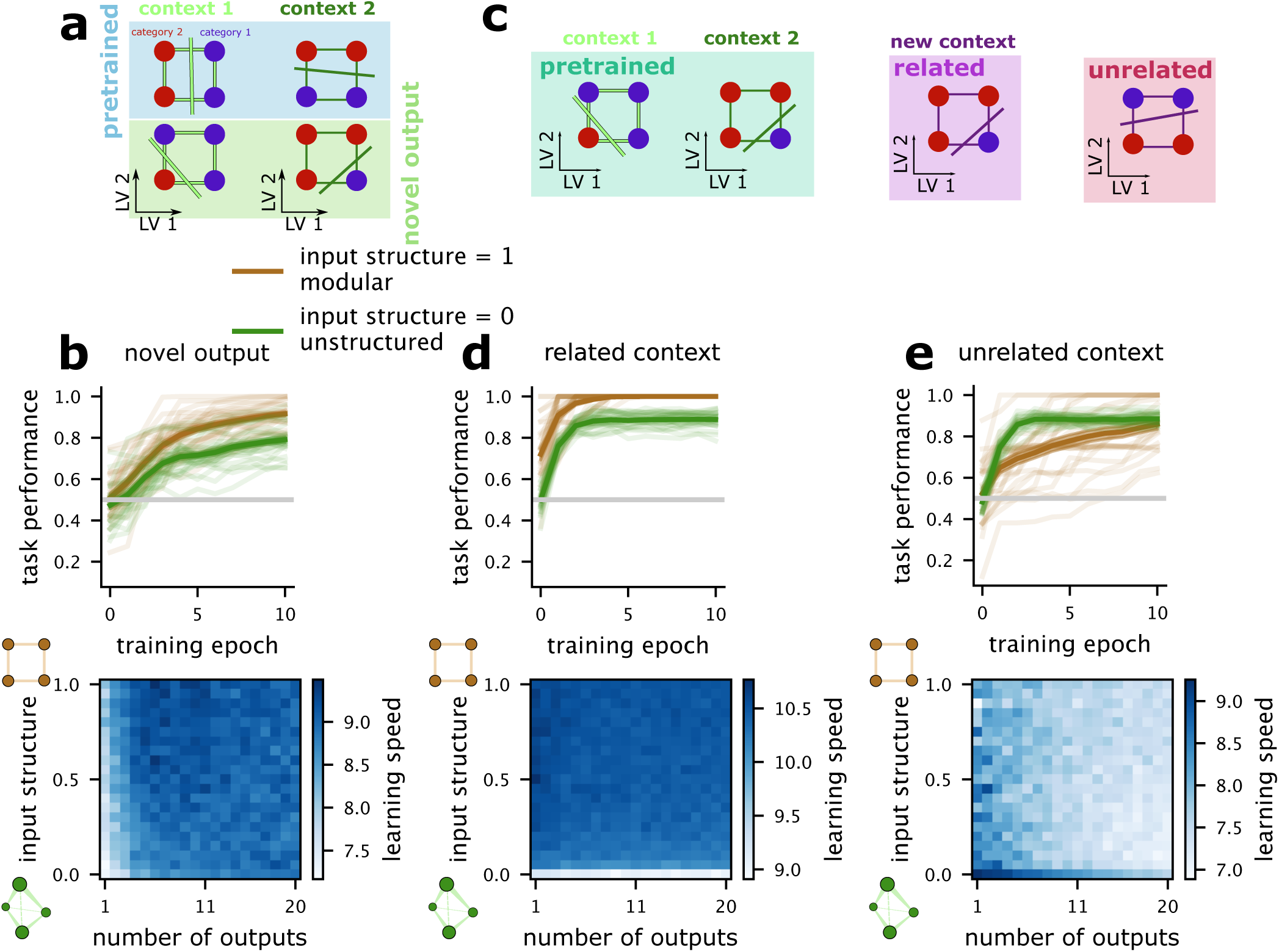
Modular representations provide rapid learning of new output dimensions and related contexts, but slower learning of unrelated contexts. **a** Schematic of the novel task learning analysis. The network is pretrained on *P* output dimensions and then trained to use a *P* + 1th output. **b** The learning trajectory of example networks pretrained to perform a contextual task with *P* = 1 output dimensions and either fully disentangled or unstructured representations (top). The learning speed of networks for the *P* + 1th output dimension, given pretraining on different numbers of output dimensions and different input geometries. **c** Schematic of the related and unrelated context learning analyses. In both cases the network is pretrained in two contexts, and must learn to perform a third context. (left) The context is related to a previously learned context, and shares all the same task boundaries as one of the two pretrained contexts. (right) The context is unrelated to either pretrained context, as has randomly selected task boundaries. **d** The same as **b** but for the related context analysis. **e** The same as **b** but for the unrelated context analysis.

Then, we investigate how the speed of learning for different kinds of novel contexts depends on both the input geometry and the number of previously trained output dimensions (fig. 4c). First, we investigate the learning of a related context (fig. 4c, “related”) in which the novel context has a different contextual input than the previously learned contexts but shares all of the task decision rules with one of the two previously learned contexts (fig. 4c, “related” and “context 2”). We show that the learning speed for a related context depends primarily on the geometry of the input representations, where higher-dimensional input representations lead to slower learning in the novel, but related context.

Finally, we investigate the learning speed of an unrelated context (fig. 4c, “unrelated”), in which the novel context has both a different contextual input than the previously learned contexts as well as different, randomly selected decision rules. In this case, the highest-dimensional input geometry provides faster learning than the low-dimensional input geometry (fig. 4e, top). However, the overall effect is more heterogeneous (fig. 4e, bottom). In particular, both decreasing the number of output dimensions and decreasing the dimensionality of input representations tend to increase learning speed.

### 2.5 More general structured tasks that give rise to modularity

We have focused on contextual tasks where each task can be decomposed into two distinct contexts and where, within each context, the required outputs are linearly separable. These tasks are naturally connected to commonly used neuroscience tasks[3, 22, 26]. Now, we generalize our analysis to tasks with distinct structure and show that modularity still emerges for nonlinear tasks that admit decomposition into common sub-tasks across different outputs. For ease of visualization, we consider these novel tasks for an input space described by three binary variables (shape: square or circle; color: green or blue; size: small or large; fig. 5a), but our findings would generalize to tasks with more input variables. We also use a more general method for visualizing and quantifying structure in the tuning of units that have learned to perform these tasks. In particular, we train artificial neural networks to perform the different kinds of tasks with many outputs (fig. 5b). Then, for each unit in the representation layer, we visualize the corresponding input weight vector (fig. 5c). This allows us to discover unexpected clusters of specialized units – or to discover that there are not distinct clusters. We quantify the amount of clustering using the density-based cluster validation score[39], which quantifies non-convex cluster structure.

**Figure 5:**
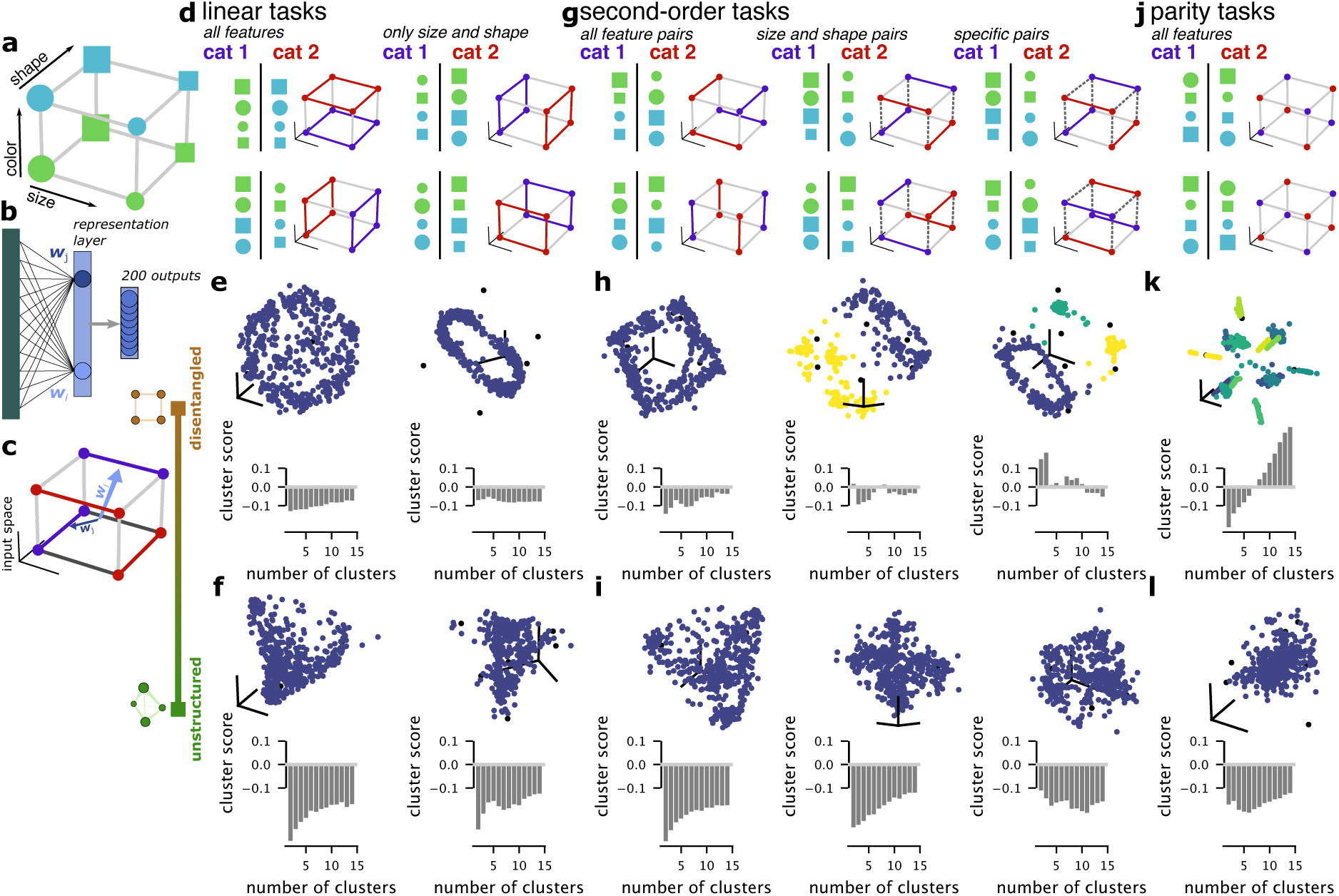
Explicit modularity emerges in networks trained to perform more general classes of tasks with disentangled inputs. **a** An example feature space, parameterized by shape (square or circle), color (blue or green), and size (small or large). **b** Networks are trained to perform different kinds of tasks with two hundred outputs. **c** We visualize selectivity in the network by visualizing the input weight vectors for all units in the representation layer. **d** Two example linear classifications of the stimulus set that either depend on all features (left) or only on size and shape (right). **e** The input weight vectors for a network trained to perform a linear task on the full space (top left) and on only two of the input dimensions (top right). The average density-based clustering validation scores of networks trained with disentangled inputs on the two task types above (bottom). **f** The same as **e** but for networks trained with unstructured inputs. **g** Example second-order tasks with different constraints. (left) A second-order task with all possible feature pairs. (middle) A second-order task with only size and shape feature pairs. (right) A second-order task with only size and shape pairs and, further, only blue circles and blue squares can be used as feature pairs to determine the output. The dashed lines indicate stimulus pairs that cannot be selected and categorized together, while the colored lines indicate stimulus pairs that were selected for a particular output. **h** The same as **e**, but for networks trained to perform the second-order tasks given in **g** with disentangled inputs. **i**. The same as **h** but for networks trained with unstructured inputs. **j** An example parity task with the two possible unique outputs. **k** The same as **e** but for networks trained to perform a parity task on disentangled inputs. **n** The same as **k** but for networks trained on unstructured inputs.

### Linear classification tasks do not give rise to modularity

First, we consider linear classification tasks in this 3D feature space. In this kind of task, each output unit must categorize the stimulus conditions according to a randomly selected hyperplane that passes through the origin of the stimulus space (so, the the classifications have the same number of stimuli in each category, but are not necessarily aligned with a particular feature axis as in the examples; fig. 5d, left, “all features”), such as to classify all shapes based on their color (fig. 5d, left, top example) or their size (fig. 5d, left, bottom example), similar to the tasks used in previous work[36]. Different colors (red and blue) correspond to the desired different values of the output. We then visualize the selectivity of the representation layer units as described above. The input weight vector corresponding to a particular representation layer unit indicates which stimulus conditions produce the strongest responses in that unit (fig. 5c). We view clusters within this space as another form of functional specialization. A set of units with input weight vectors pointing in a consistent direction within this space will all respond to the same set of stimuli. A set of units with weight vectors pointing in a different direction will respond to distinct sets of stimuli. These different sub-populations of units in the representation layer will decompose the task into distinct sub-tasks, just as with the contextual tasks from before.

For networks with disentangled input representations trained to perform linear tasks, the tuning covers the full space of inputs and does not have distinct clusters (fig. 5e, top left). Unlike our contextual task from before, the different outputs here do not share any structure (such as a shared context variable). The full task is also already linearly separable in the input space. We confirm the absence of distinct clusters through spectral clustering in the weight space. We choose spectral clustering because, even for linear tasks, the tuning is highly structured. For instance, nearly all units are well away from the origin, indicating they have significant tuning for the stimuli. In many traditional clustering approaches, this would be inconsistent with them being a single cluster, since the cluster is non-convex. Spectral clustering recognizes this structure as a single connected component and, therefore, a single cluster. We compute the density-based clustering score[39] for 2 to 15 clusters. If no number of clusters has a positive validation score, then we consider the tuning best fit by a single cluster. This is the case for the linear task with disentangled inputs described above (fig. 5e, bottom left), where stimuli are classified according to a random linear hyperplane in the input space. We repeat the analysis for the same kind of linear task but given unstructured, high-dimensional inputs (fig. 5f), which also yields tuning without significant clusters. Next, we consider a variant on this linear task, where the output depends only on a subset of the three variables (e.g., only on size and shape; fig. 5d, right, “only size and shape”), for instance classifying all large from small stimuli (fig. 5d, right, top example) or circles from squares (fig. 5d, right, bottom example). Networks trained to perform a task with this output structure and disentangled inputs develop tuning in a ring in the input space, capturing the variance of these two variables, but not of the third (fig. 5e, top right). In this case, too, our clustering analysis indicates that there is only a single cluster (fig. 5e, bottom right), though, again, that cluster is highly non-convex. Networks trained to perform this task with unstructured inputs (fig. 5f, right) also do not develop distinct clusters.

### Second order nonlinear tasks that do not give rise to modularity

Next, we turn to nonlinear tasks that depend on pairs of feature values, which we term second-order tasks. Here, the target output for a particular stimulus is determined by a pair of its features (for instance, its size and shape), rather than by a single feature as with the linear task (for instance, its color, as in fig. 5d, top left). A possible output would have green squares and small blue shapes in one category (fig. 5g, top left, “all feature pairs,” “cat 1”) with green circles and large blue shapes in the other category (fig. 5g, top left, “cat 2”). In this case, any pair of features could be chosen – so, a different output could have large squares and small circles in one category with large circles and small squares in the other (fig. 5g, bottom left). As stimuli are assigned to a particular category as an adjacent pair, we have represented this structure in the visualization of the output, where the line connecting two stimuli is colored according to the category of that pair (fig. 5g). Notice that the colored lines can be different for different outputs.

Networks trained to perform this kind of task develop highly structured tuning that reflects the input space and the dependence of the task on pairs of stimuli (fig. 5h, top left), but that does not split into distinct clusters for either disentangled (fig. 5h, bottom left) or unstructured inputs (fig.Latent variable 5i, left).

### Second order nonlinear tasks produce modularity

We explore two other variants on second order tasks, constructed by excluding different feature pairs from shaping the target output. In the first case, only size and shape pairs can be chosen to determine the target output (fig. 5g, middle, “size and shape pairs”). This structure replicates our contextual tasks from before, since now color acts as a contextual variable where, stimuli of one color are classified according to one linear boundary but stimuli of the other color are classified according to a different linear boundary (fig. 5g, middle). This structure can be imposed on the second-order tasks described above by excluding certain stimulus pairs from selection to be in a particular category (fig. 5g, middle, dashed lines). Importantly, this structure is shared by all the output units (the dashed lines are the same), analogously to the case of context-dependent tasks.

Networks trained to perform this kind of task have the selectivity expected from our previous results, where networks trained with disentangled inputs develop distinct clusters. One cluster has units that are selective for stimuli from one context (or with one color, fig. 5h, top middle, yellow points) and the other has units that are selective for stimuli from the other context (or with the other color; fig. 5h, top middle, navy points). Despite the non-convexity of these context-dependent clusters, we find significant clustering scores (fig. 5h, bottom middle) for two and eight clusters, as expected from our previous results. We also find that networks trained with unstructured inputs do not show significant clustering (fig. 5i, middle), which is also consistent with our prior results – though we do expect these networks to have implicit modularity.

In a second variant on second-order tasks, we restrict the feature pairs that can be selected even further by excluding large blue shapes and small blue shapes from shaping the target output (fig. 5g, right, dashed lines). This means that blue circles and blue squares will always shape the target output as pairs (fig. 5g, right). In this case, networks trained with disentangled inputs develop three clusters in tuning (fig. 5h, right). One diverse cluster with units responding to combinations of green shapes, and two smaller clusters, in which units respond either only to blue circles and or only to blue squares (fig. 5h, right). This task structure can also be viewed as equivalent to the contextual task case, where the outputs in one of the two contexts (when the color is blue) always classify the stimuli according to a single feature (their shape), while the outputs in the other context are unconstrained. It is this consistent structure across the outputs that shapes the clusters in tuning. Networks trained with unstructured inputs, in contrast, develop no reliable clusters (fig. 5i, right).

### Highly nonlinear tasks produce very selective modules

Finally, we explore the tuning developed for a third kind of highly nonlinear task, that we term parity tasks. In this task, each stimulus has its output determined by all three of its features. In particular, if each feature value is represented as a one (e.g., small, green, circle) or a zero (e.g., large, blue, square), then the stimulus will be in category 1 if the sum of those feature values is even (fig. 5j, “cat 1”) and in category 2 if the sum of those feature values is odd (fig. 5j, “cat 2”). For a three-dimensional stimulus space, there are only two unique parity task outputs, one mapping odd stimuli to category 1 and the other mapping odd stimuli to category 2 (and vice versa for even stimuli; fig. 5j, top and bottom). Networks trained to perform this parity task with disentangled inputs develop highly specific tuning in their representation layer, where each stimulus is represented by an almost unique set of units (fig. 5k). In contrast, networks trained with unstructured inputs do not develop clustered tuning (fig. 5l).

Our results in this section generalize our findings from tasks with a contextual structure to tasks with a diversity of structure. In this more general context, we find an analogous set of results: only networks trained from disentangled input representations develop distinct, functionally specialized sub-populations of units in their representation layer. Networks trained on unstructured inputs never develop these sub-populations. This makes sense because, for a completely unstructured input representation, all of these tasks are essentially equivalent: they are all random ways of labeling a set of uncorrelated points. We also find that these specialized sub-populations only emerge when the task itself can be decomposed into multiple linearly separable sub-tasks that are consistent across outputs and concern distinct sets of points (fig. 5h, middle and right, as well as fig. 5k).

## 3 Discussion

We have shown how the learned representational geometry of an artificial neural network trained to perform contextual behavior depends on the geometry of its input and the number of task output dimensions. These learned geometries have been observed in the brain, and range from strikingly modular representations where individual units are only active in a specific context to completely unstructured, high-dimensional representations, which enable any task to be performed. We provide intuitive explanations for the transitions between these different regimes. We also explained the computational advantage of modularity in contextual tasks: context dependence almost always introduces non-linear separability, and the different modules reflect a possible decomposition of the full context-dependent task into subtasks (a subset of conditions) that are linearly separable. We have also demonstrated how our results change under different learning regimes (e.g., rich or lazy learning[26, 40]; see *Learned representations depend on the learning regime* in *Supplement* and fig. S1). Then, we show that these different learned representations have distinct computational benefits, enabling different forms of generalization. These results demonstrate that low- to moderate-dimensional input representations to networks trained to use a handful of output dimensions yield benefits for both learning novel output dimensions and for generalizing to unseen stimuli. We demonstrate how these learned representations shape the learning of future contexts – and demonstrate differential benefits for learning related and unrelated contexts. Finally, we defined a more general set of tasks and showed that the intuitions gained from the contextual task case apply also for more complex, arbitrary tasks.

### 3.1 Predictions for experimental data

Our work makes several distinct predictions for experimental data. First, we predict that the representation of contextual and decision-related variables prior to training or in preceding brain regions will strongly influence how training shapes the representational geometry in the animal. The representations generated before training are likely to mirror the structure of the input data. For instance, if we assume that the weights connecting the input to the neurons in the representations we are examining are initially random and uncorrelated with the inputs, we can conclude that the geometry of the input is almost perfectly recreated in the initial representation when the neurons are linear[41, 42]. When the neurons are nonlinear, the geometry is only approximately replicated, but it is often surprisingly similar, as the ranking of the distances is preserved[43, 44]. In particular, if those initial representations are low-dimensional and disentangled, then our framework predicts that modular representations will be more likely to emerge. In contrast, if those initial representations are high-dimensional and unstructured, then our framework predicts that the representation will not become explicitly modular – that is, with specialized sub-populations of neurons for different task conditions – but that it may become implicitly modular depending on the nature of the task and the dimensionality of the output. The point at which we expect a transition between these two regimes is also specified by our theory, and depends on the complexity of the stimulus feature space (see *Explicit modularity emerges when the disentangled component of the input provides faster learning* in *Supplement*) as well as on the learning regime (see *Learned representations depend on the learning regime* in *Supplement*). However, not all representations in the brain need to change to support task performance – so, for instance, we do not expect large changes in earlier sensory regions, where task variables may not be linearly separable. These predictions emphasize the importance of longitudinal and multi-region recordings in neural data.

#### Predictions for learning

Our framework predicts that these different situations will lead to different levels of robustness to changes to the experimental framework. We expect unstructured representations to be associated with highly brittle behavior and a failure to generalize to subtle changes in the experimental setup. In contrast, we expect modular representations to be associated with a greater robustness to irrelevant changes (e.g. changes in latent variables that are not relevant for performing the task). This set of predictions could provide an explanation for what have remained largely anecdotal findings of brittleness to certain kinds of changes in experiments – for instance, animals are often found to fail to generalize to even subtle changes in experiments yet regularly generalize in more naturalistic conditions. This could be because the variables that are relevant to more naturalistic behavior are low-dimensional and disentangled (as predicted by previous work [36]) while representations of various experimental variables are more high-dimensional. In addition, due to different training histories, different animals could have different input, or initial, representational geometries when they are being trained to perform the same task. These different initial geometries could lead to differences in representational geometry once the animal is fully trained; such differences have previously been shown to correlate with different behavioral strategies[37].

Finally, our framework predicts that different learned geometries will give rise to different learning speeds for novel behaviors – and that this effect will depend on whether the behavior is related or not to previously learned behaviors. This prediction can be tested in experiments where an animal sequentially learns a series of interrelated tasks. In particular, if we find that the representational geometry underlying task performance is either explicitly or implicitly modular, then we expect the animal to rapidly learn related tasks but struggle to learn unrelated tasks; in contrast, if we find that the representation geometry is highly unstructured, then we expect that the animal will learn related and unrelated tasks at the same rate.

### 3.2 Connection to prior work

Our work significantly extends previous work focused on neural networks trained to perform contextual behavior[24–26, 32]. In particular, previous work has only considered low-dimensional input and single output (i.e., single task) conditions – while we consider the full interaction space between these two parameters. We believe that considering a much wider variety of input geometries is essential precisely because such a diversity of representational geometries has been found in experimental work – ranging from low-dimensional face representations[33, 34] to high-dimensional representations of cognitive variables in frontal cortex[21, 31]. This work provides a unified framework for understanding learning from these different starting points – and indicates that such diversity may be essential to explain the wide diversity of representational geometries reported to underlie contextual behavior in the literature[3, 21, 22]. Similarly, we believe that considering an increased number of output dimensions used within each context is important because natural behavior is often richer than a single binary decision – instead, multiple different decisions are made about a stimulus at once, and these decisions produce multi-dimensional output that has so far remained relatively understudied in experimental neuroscience.

#### Modularity and compositionality

One aspect of previous work that we have not extensively characterized here is the notion of compositionality[24, 25, 45] in the learned representations. In prior recurrent neural network studies, they found that, in some cases, the same neural subspace would be used to represent the continuous value of a particular stimulus feature across different contexts. The networks trained to perform a contextual task that is related to one of several previously learned tasks are compositional. In this case, they re-use the same subspace used to perform the original task to perform the new task, and this re-use likely explains the faster learning for explicitly modular, as opposed to implicitly modular or unstructured representations.

Our work also only focuses on one form of contextual decision-making: when different contexts are associated with different task rules[3, 10, 22, 23, 46]. However, different contexts can also be associated with different thresholds for behavioral response[47] or biases in the stimuli that are shown[48]. Elucidating the computational principles that shape neural representations in these cases will require further work. However, it is possible that the initial representational geometry will be crucial in these cases as well, since the interaction between representations of decision variables and the representation of the context cue will also shape the kinds of solutions learned in artificial neural networks. Thus, the principle of an initial representation shaping the learned representation is likely to still be important in this case.

#### Modularity and categorical representations

The definition of explicit and implicit modularity that we use here is related to, but not the same as, the categorical and non-categorical selectivity discussed in related work[6, 49]. Categorical selectivity in a neural population is defined by the existence of clusters of neurons with similar response profiles to the full set of conditions in an experiment. Our analysis of clustering in the weight space of neural networks trained to perform a diversity of tasks (fig. 5) is related to the approach used in this work. There, we showed that distinct clusters existed only for networks trained with low-dimensional, disentangled input representations. These clusters also fit our definition of modularity, since the different clusters of units responded to different subsets of the full set of stimulus conditions. Thus, in this setting, explicit modularity implies categorical selectivity – however, the reverse is not necessarily true. In particular, we define explicit modularity as two or more sets of neurons that representation distinct stimulus conditions. Categorical selectivity does not require these distinct clusters of neurons to represent distinct sets of stimulus conditions; instead, two well-separated clusters of neurons could respond to the same set of stimulus conditions, but one cluster responds strongly and the other weakly. In contrast, implicit modularity does not necessarily imply either categorical or non-categorical selectivity, though the lower-dimensional structure in neural responses associated with implicit modularity in our definition would necessarily manifest as low-dimensional structure in the condition space used in[6]. In our clustering analysis, we also chose an approach meant to quantify gaps in selectivity even when all selectivity is highly structured. This is because, in our models, most units end up tuned to some subset of the stimulus conditions after training, but the same is not true of neurons in the brain. Further work is necessary to provide a more comprehensive link between these two frameworks for thinking about structure in neural representations.

## Conclusions

Overall, our results provide an understanding of why different representational structure emerges in different situations. We argue that knowledge of the geometry of the initial representations of the task-relevant variables, which is likely to reflect the structure of the inputs, is essential for predicting the learned structure in the representations. This core prediction has already been validated in some experimental datasets, and underlines the importance of collecting data that allows the characterization of this initial geometry. Further, our work makes predictions for how these learned representations will, in turn, shape the learning of future behaviors and the ability to generalize to novel stimuli.

## Acknowledgments

We are grateful to Matteo Alleman, Samuel Lippl, and members of the Center for Theoretical Neuroscience for useful discussions. We are grateful to Allison Ong and Ciela Sophia Chavez-Gilbride for administrative support. This work was supported by the following grants and foundations: NIH NINDS K99NS138578 (WJJ), Simons Foundation 542983SPI, Gatsby Charitable Foundation GAT3708, the Kavli Foundation, and the Swartz Foundation. We acknowledge computing resources from Columbia University’s Shared Research Computing Facility project, which is supported by NIH Research Facility Improvement Grant 1G20RR030893-01, and associated funds from the New York State Empire State Development, Division of Science Technology and Innovation (NYSTAR) Contract C090171, both awarded April 15, 2010.

## Author contributions

WJJ and SF designed the framework. WJJ performed the simulations and analyzed the models. WJJ made the figures. WJJ and SF wrote and edited the paper.

## M1 Methods

### M1 Latent variables

The input consists of *L* = *D*+*I*+*C* binary latent variables, where *D* are the decision-related variables (i.e., variables that are used in classification tasks), *I* are irrelevant variables (i.e., variables that do not influence the target output), and *C* are context variables. Unless otherwise noted, there are *D* = 3 decision variables, *I* = 4 irrelevant variables, and *C* = 2 context variables. The decision and irrelevant variables are each independently sampled from a binomial distribution with *p* = .5. In contrast, the context variables are constrained to be one-hot across the *C* context dimensions, and are sampled with equal probability of each context being active. Instead of taking on the values 0 and 1 all variables are then remapped to take on the values −1 and 1 for notational convenience.

### M2 Input model

Within the input model, all *L* latent variables are treated identically. We consider a spectrum of input geometries. Low-dimensional, disentangled representations are at one end of the spectrum and for samples *x* ∼ *X* where *X* is the distribution of latent variables (see *Latent variables* in *Methods*), the representation of the stimuli in the disentangled input model are given by,

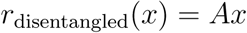

where *A* is a *N*_input_ × *L* matrix, where *N*_input_ is the number of units in the input model and *L* is the number of latent variables. The rows of *A* are chosen to be orthogonal to each other with *N*_input_ *> L* and the magnitude of the elements of *A* are chosen so that the sum of variance across *r*_disentangled_ is 1.

High-dimensional, unstructured representations are at the other end of the spectrum, they are given by,

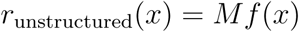

where *f* (*x*) produces vectors with length 2*^L^*according to,

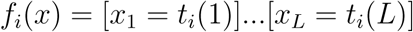

for *i* ∈ [1*, …, n^L^*] where [*x* = *y*] is the indicator function which is 1 when the *x* and *y* are the same and 0 otherwise, as used in prior work[12, 13, 21], and *t_i_* is either −1 or 1. The matrix *M* is *N*_input_ × 2*^L^* and *N*_input_ *>* 2*^L^* with all rows of *M* chosen to be orthogonal with scaling chosen so that the sum of variance across *r*_unstructured_ is 1. Thus, each unique stimulus described by the latent variables *L* is represented in an independent dimension.

Finally, to produce a full spectrum of codes, we linearly interpolate between these two extreme cases, such that,

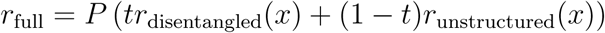

where *t* ∈ [0, 1] and *P* provides an overall scaling of the representation. Throughout the manuscript *P* = 1.

### M3 Contextual tasks

The contextual tasks are given by,

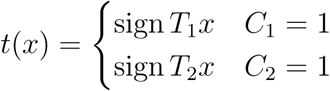

where *T_i_* are randomly sampled unit-length, *L*-dimensional vectors. Each *T_i_* is sampled independently. Note that only one *C_i_* can be 1 for any stimulus *x*, so the cases are mutually exclusive.

### M4 Contextual multi-tasking model

The contextual multi-tasking model is a feedforward network with a single hidden layer. The inputs and target outputs are described above. The main parameters of the model are:

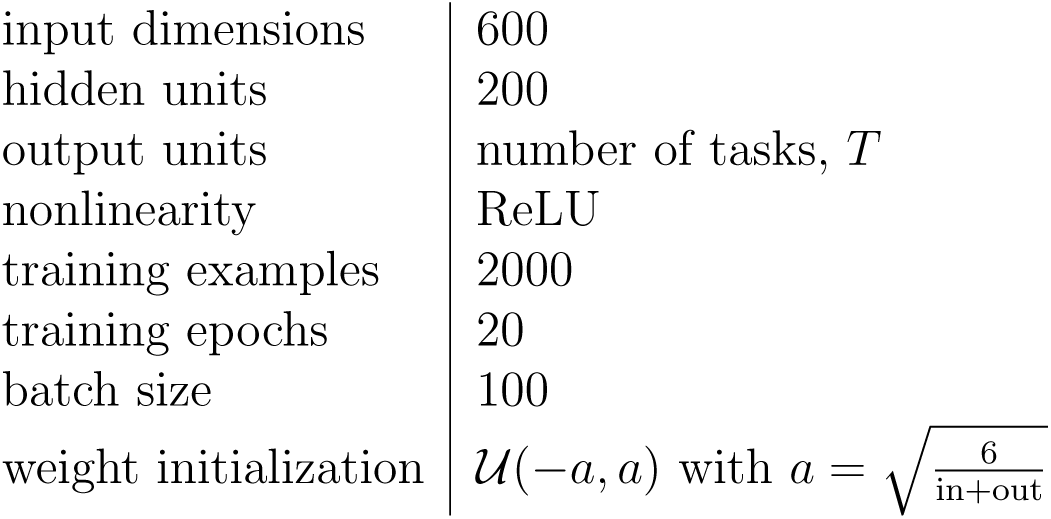

|  |  |
| --- | --- |
| input dimensions | 600 |
| hidden units | 200 |
| output units | number of tasks, $T$ |
| nonlinearity | ReLU |
| training examples | 2000 |
| training epochs | 20 |
| batch size | 100 |
| weight initialization | $\mathcal{U}(-a, a)$ with $a = \sqrt{\frac{6}{\text{in}+\text{out}}}$ |

The network is trained with the optimizer Adam and a learning rate of 1e-3.

### M5 Dimensionality of unstructured solutions

The unstructured representation that we consider in the manuscript is primarily the maximum-dimensionality unstructured representation, where units have conjunctive responses for combinations of all *L* latent variables. This representation has dimensionality that scales exponentially with the number of latent variables, as dim(*x*) = 2*^L^*. While such high-dimensional interactions are necessary for a linear decoder to be able to learn any binary labeling of the stimuli, there are lower-dimensional but still unstructured representations that would allow a linear decoder to learn additional classes of labelings.

These other classes of unstructured representations can be described by the order of their interactions. A representation with order *O* = *L* is the maximally unstructured representation from before. A representation with order 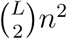 interaction terms (e.g., an interaction between latent variable *i* and *j*, [*x_i_* = 0] [*x_j_* = 0] where the brackets represent the indicator function as in *Input model* in *Methods*) and where *n* is the number of values that each latent variable can take on. In general, an unstructured representation with order *O* tuning will have 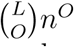 interaction terms. Importantly our measure of subspace specialization is robust to unstructured representations of different orders – that is, an unstructured representation of any order will still have subspace specialization equal to zero.

An *O* = 2 unstructured representation can be used to solve any contextual task. Thus, we calculate the dimensionality (described by the participation ratio) of this class of unstructured representation. The participation ratio can be written as [50],

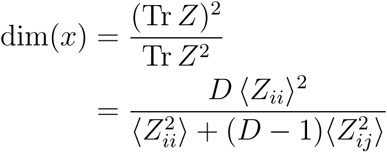

where *D* is the number of dimensions in the input and *Z* is the *D* × *D* input dimension covariance matrix, across all stimuli.

We calculate,

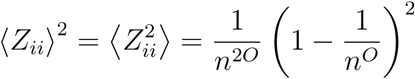

And

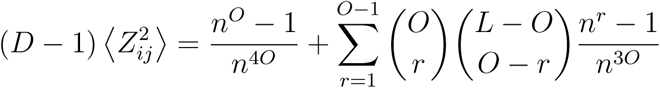

So, putting it together, we obtain,

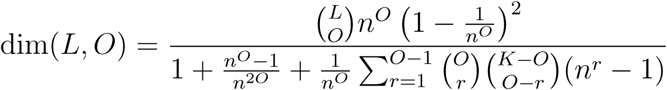

where *L* is the number of latent variables and *O* is the order of the interactions. For *O* = 2, we find that,

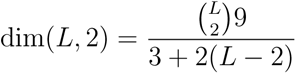

which will grow proportionally to *L* for large *L*.

### M6 Clustering analysis

We assess whether the units in the hidden layer of the complex task network are best explained by discrete clusters in the space of average activity within each context. To do so, we sample 1000 stimuli from each context and take the average activity of each unit within each context. This produces a matrix of dimensions *µ_C_* = *N*_hidden_ × *C*. Then, we perform Gaussian mixture model clustering on this matrix, where each unit is assigned to one of *K* discrete clusters. We perform this procedure for *K* = 1 to *K* = 5. Then, we evaluate the goodness of fit of each clustering with the Bayesian information criterion (BIC)[51], and choose the clustering with the highest BIC. We use these clusters to color the points in fig. 2d and similar plots. Where only one color is present, this means a single cluster provided the best fit to the data.

### M7 Contextual fraction

We define the context fraction in the same mean activity space described above for the clustering analysis. Here, after computing *µ_C_*, we then count the rows of the matrix that have mean activity *<* 0.01 in all contexts except one, this count is then divided by the number of units in the hidden layer.

### M8 Subspace specialization

We define subspace specialization to capture whether the population activity in the hidden layer of the complex task network has low-dimensional structure related to context or if it is simply high-dimensional and unstructured. The measure is designed to be 0 if there is no unique low-dimensional structure related to context and positive otherwise. The measure is based on the alignment index, which has been introduced previously in [35]. The alignment index quantifies the amount of overlap between the subspaces described by two sets of basis vectors, *U*_1_ and *U*_2_, which are both *N* × *P_i_* matrices where *P_i_* is the number of basis vectors. The alignment index for *U*_1_ and *U*_2_ is,

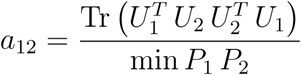

A value of 1 indicates that the two sets basis vectors span precisely the same subspace, while a value of 0 indicates that they span non-overlapping subspaces of the full population activity space.

To compute our subspace specialization measure, we compute *D* + *I* non-context alignment indices and 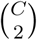 context-related alignment indices. For each latent variable (or context pair), we sample 1000 stimuli and then split into two subsets, where the value of the latent variable is zero (or the first context is active) in one subset and where the value of the latent variable is 1 (or the second context is active) in the other subset. Then, we perform PCA separately on the two subsets, and keep all dimensions for each that explain more than 10*^−^*^10^ of the variance. This gives us our sets of basis vectors, *U*_1_ and *U*_2_. We take the alignment index between these sets of vectors, as defined above. Now, we have *D* + *I* overlap measurements for non-context variables, *o*_noncontext_, and 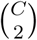 overlap measurements for context pairs, *O*_context_. We summarize these measurements into a single subspace specialization measure by taking the average of each group, *O̅*_noncontext_ and *O̅*_context_, followed by the rectified log-ratio:

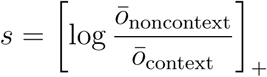

for the subspace specialization index. This index will be zero when the alignment for the context pairs and non-context variables is the same. It will also be zero (due to rectification) when the context variables have greater alignment than the non-context variables. However, it will be positive when the context variables have lower alignment than the non-context variables. This indicates that the activity in different contexts is confined to subspaces that are distinct relative to that of other variables encoded by the same system.

### M9 Alternate decompositions

We want to calculate the probability that a the decomposition along a given dimension is eliminated (i.e., made so that decomposing along that dimension does not yield linearly separable sub-tasks) by a randomly selected task. We simplify our tasks for this calculation by assuming that they are all aligned with a single decision-relevant latent variable. We do not make this assumption when testing our prediction.

For a given task and with *D* relevant latent variables, there are three possible situations. With probability 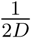, the same decision-related variable and direction along that variable will be selected. This whole task is already linearly separable and it does not eliminate any dimensions. With probability 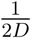, the same decision-related variable but different directions will be selected. This eliminates all variables as possible decompositions except for the selected variable. With probability 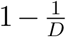, two different decision-related variables will be selected. This eliminates all variables except for these two variables.

To put these together, we compute the probability that a particular variable is not eliminated after a new task is added. This is the sum of the probability that this particular variable is chosen at least once or it is not chosen at all, but the the same variable and direction is chosen. The probability that it is chosen at least once is,

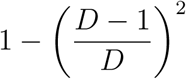

and the probability that the same other variable is chosen twice in the same direction is,

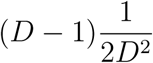

So, together, the probability that a given variable is not eliminated after a single task is selected is,

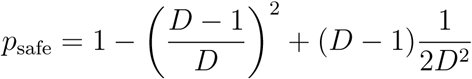

and, because each task is selected independently, the probability that it eliminated after the selection of *T* tasks is,

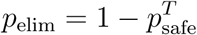

where *T* is the number of tasks. Finally, the expected number of alternate decompositions is then

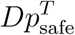

### M10 Dimensionality of the required output

We compute the participation ratio of the required output, *x* averaged across all stimuli. To approach this, we employ the expression for the participation ratio developed in [50],

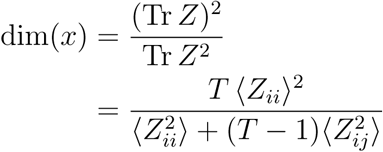

where *T* is the number of tasks and *Z* is the *T* × *T* task covariance matrix, across samples from all contexts. The diagonal elements of the covariance matrix *Z_ii_* are all defined to be 1, so 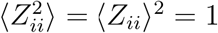. The off-diagonal elements require calculation. So, we want to calculate 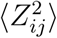 ⟩ for

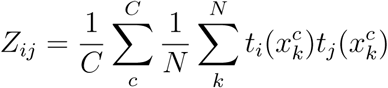

where *C* is the number of contexts, *N* is the number of samples taken from each context, and 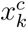 is the *k*-th sample from context *c*.

First, we simplify our tasks so that they are aligned with the decision-related variables in each context. We do not make this assumption elsewhere in the paper, nor when verifying the predictions that arise from this calculation. In this simplified setting, the average product within each context can take on three different values: 1 when tasks *i* and *j* are the same (i.e., they are aligned with the same decision variable *D* and pointing in the same direction), and this happens with probability 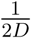; 0 when the tasks are orthogonal (i.e., aligned with different decision variables), and this happens with probability 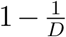; and −1 when the tasks are anti-parallel (i.e., aligned with the same decision variable, but pointing in different directions), and this happens with probability 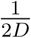.

So, we want to calculate,

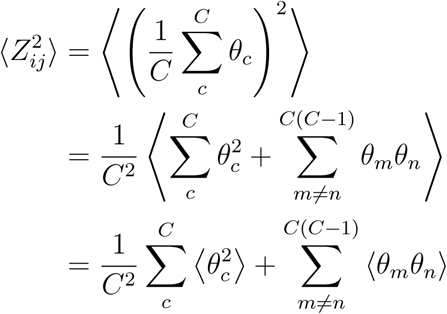

Where

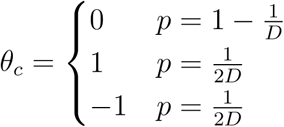

So,

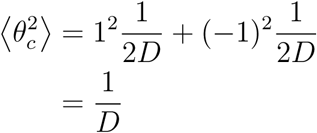

and

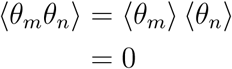

which yields

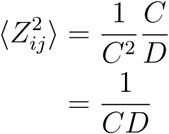

Putting everything together, we find

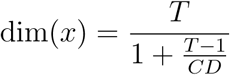

### M11 Cross-condition generalization performance

To compute the cross-condition generalization performance (CCGP) within a particular context, we first select one decision-relevant latent variable for decoding. Then, we train a decoder to decode the value of that variable given 8 stimulus representations sampled from the same context and where all other decision-related variables have a fixed value. Finally, we test the same decoder on 8 stimulus representations sampled from the same context but where the other decision-related values never have the their fixed value from before. We repeat this 10 times and report the average performance of this decoder as the CCGP. All samples have the same noise level as used in network training.

### M12 Shattering dimensionality

To compute the shattering dimensionality within each context, we select all balanced dichotomies of the stimuli based on their decision-relevant features. Then, we train a linear decoder to perform that classification task using 1000 samples and report its performance on 1000 test samples with different noise. All samples have the same noise level as used in network training.

## S1 Supplement

### S1 Learned representations depend on the learning regime

We have focused on a rich learning regime in the rest of the paper. In this regime, the representations change appreciably as the network is trained. Intuitively, modular representations are less likely to emerge in the lazy learning regime, as the representation in the network will not change as much over training. To give a sense of how our results depend on the learning regime, we train three sets of networks with different weight initialization statistics. The first set is close to the one investigated in most of the paper. This rich regime network is has its weights initialized following *w_i_j* ∼ N(0*, σ*) with *σ* = .001 (fig. S1a). The second set has larger initializations and is closer to the lazy regime (*σ* = .02; fig. S1b). The third set is trained in an even lazier regime (*σ* = .8; fig. S1c). The change of learning regime shifts the threshold input geometry where the network transitions from learning explicitly modular representations to learning unstructured or implicitly modular representations; however the results are otherwise qualitatively similar.

### S2 Intuition about the computational advantage of modularity

Consider the examples in fig. S2a,b. We assume that there are two contexts with two decision-relevant variables (so, 8 total stimuli), and we start by considering one task that separates the 8 stimuli into two groups of 4 stimuli each (e.g., fig. S2a). To further simplify the argument, each task is constrained to be aligned with one of the relevant latent variables. While we assume that the contextual task is linearly separable within each context, the full task across both contexts will not be linearly separable in most cases – in fact, the full task will only be linearly separable if the task selected in both contexts is exactly the same (see *Alternate decompositions* in *Methods* for details). This is the problem that the output dimensions has to solve (separate the 8 points into two groups according to their color in fig. S2a,b).

The modular representation provides a simple solution to this problem: some of the units in the intermediate layer could be active only in context 1 (unit 1 and 2 in fig. S2a), and the others in context 2 (unit 3 and 4). This will always yield a representation that makes the full task linearly separable, since we assumed that the task is linearly separable in each context. Then, in context 1, the output unit tunes the weights between it and the active neurons. The weights from the other units are irrelevant because these other units are inactive. The same procedure can be repeated in context 2. In other words, by dividing the 8 points into two groups of four points, the network is able to ‘handle’ each group with separate sets of units and yield a representation that makes the full task linearly separable for the output unit.

This modular decomposition based on context is clearly a solution (fig. S2a, b, “context decomposition”), but it is typically not the only solution. Indeed, there are other ways of decomposing the problem and dividing the 8 points into two groups (fig. S2a,b, “non-context decomposition” and fig. S2c). However, we show that, when the number of output dimensions increases, the number of non-context decompositions that make all outputs linearly separable shrinks rapidly (fig. S2d, left, and see *Alternate decompositions* in *Methods*). In contrast, the context decomposition is guaranteed to make all output dimensions linearly separable for any number of outputs. The modular solution then has a computational advantage over these alternative decompositions. For simplicity we discussed a solution that is explicitly modular, but the same arguments would apply to implicit modularity. Indeed, an implicitly modular solution is just an explicit one that is rotated, and the same logic of non-interference between learning in different contexts applies in this rotated space as well.

**Figure S1:**
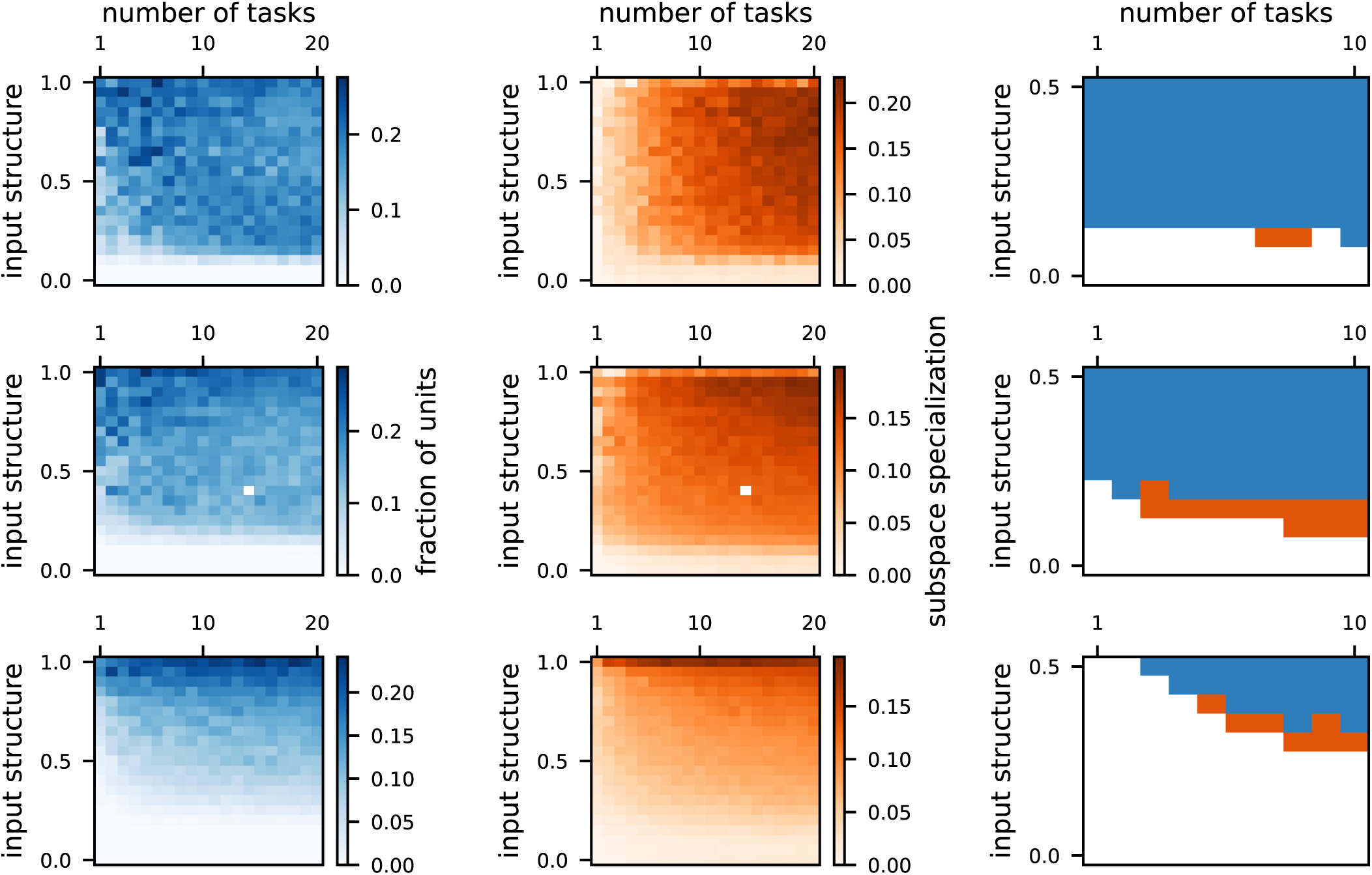
The learning regime of the network influences the learned representations. **a** Results for networks trained with small weight initialization (*σ* = .001). (left) The explicit modularity. (middle) The implicit modularity. (right) Threshold transition plot, focusing on a subset of the full parameter spacer. **b** The same as a but for larger weight initialization (*σ* = .02). **c** The same as a but for an even larger weight initialization (*σ* = .8).

If this understanding is correct, then we expect that networks performing only a single task may have lower subspace specialization – and less modular structure – than networks performing contextual tasks with multiple output dimensions. To test this hypothesis, we calculate the average number of non-context decompositions that yield a linearly separable representation as a function of both the number of output dimensions and the number of relevant latent variables. To approach this, we view each task as eliminating certain decision-related variables but not others as candidates for decomposition, depending on how the two linear sub-tasks relate to each other (fig. S2c, and see *Alternate decompositions* in *Methods* for details). We find that more decision-related variables yield fewer possible non-context decompositions (fig. S2d, left). Thus, we also expect more decision-related variables to be associated with greater subspace specialization for the same number of output dimensions, and our simulations confirm both of these predictions (fig. S2d, right).

However, we do not expect unstructured input geometries to be constrained in the same way. In this case, the full contextual task is already linearly separable in the input – and therefore decomposition into sub-tasks is not necessary. However, the contextual task is still lower-dimensional than the input geometry, so we expect our network to find a solution with dimensionality in between the dimensionality of the input and of the required output (fig. S2e)[1]. We expect this low-dimensional solution to inherit its structure from the structure of the required output. Thus, we expect the subspace specialization in the representation to follow the subspace specialization in the required output. To test this, we calculate the subspace specialization for the output for different numbers of decision-related variables (fig. S2f). Then, we show that this pattern of emergence matches the pattern that we see in the representation layer of our trained networks (fig. S2g). Interestingly, this pattern is the opposite of what we found for the disentangled inputs. For disentangled inputs, subspace specialization emerges more quickly for more decision-related variables; in contrast, for unstructured inputs, subspace specialization emerges more quickly for fewer decision-related variables. This further illustrates how changes to the input geometry alter learning dynamics in the network. However, the overall cause of the emergence of subspace specialization remains the same: The presence of low-dimensional structure within the tasks that the network exploits in different ways for different input geometries.

### S3 Explicit modularity emerges when the disentangled component of the input provides faster learning

The input geometries used to train our models (fig. S3a) can be decomposed into two components: a structured, disentangled component (fig. S3b, brown) and an unstructured component (fig. S3b, green). The input structure is varied by changing the relative strength of these two components while keeping the overall strength of the full representation constant (see *Input model* in *Methods* for details). So, an input geometry with high structure has a large disentangled component and a relatively small unstructured component (fig. S3b, top); in contrast, an input geometry with low structure has a small disentangled component and a large unstructured component (fig. S3b, bottom).

We hypothesize that, in most cases, learning is dominated by one of these of two components. This follows from previous work that shows winner-take-all dynamics for learning in artificial neural networks[2], where possible solutions with a small initial advantage in learning speed come to dominate the learned solution. This prior work shows that for disentangled inputs and a network trained to perform a single contextual task, the explicitly modular solution provides the fastest learning – and dominates the resulting hidden layer representation[2]. Here, we hypothesize that our networks have explicit modularity precisely when learning from this disentangled component of the input is faster than learning from the unstructured component of the input. To test this, we train models on the decomposed inputs, where one model is trained only on the disentangled component of an input geometry with a given level of structure (fig. S3c, yellow line) and a second model is trained only on the unstructured component of the geometry (fig. S3c, green line). We then take the difference between the average loss of these models across training (fig. S3c, “Δ learning” and right) and repeat this procedure for many different input structures: for high input structure, the disentangled component will be large in magnitude and the unstructured component small; and vice versa for low input structure. As the input structure changes, the green and brown curves change because the magnitude of the representation they are learning from changes in size. In particular, for high input structure (fig. S3c, top), the brown curve reflects learning from a large disentangled component and the green curve reflects learning from a small unstructured component.

**Figure S2:**
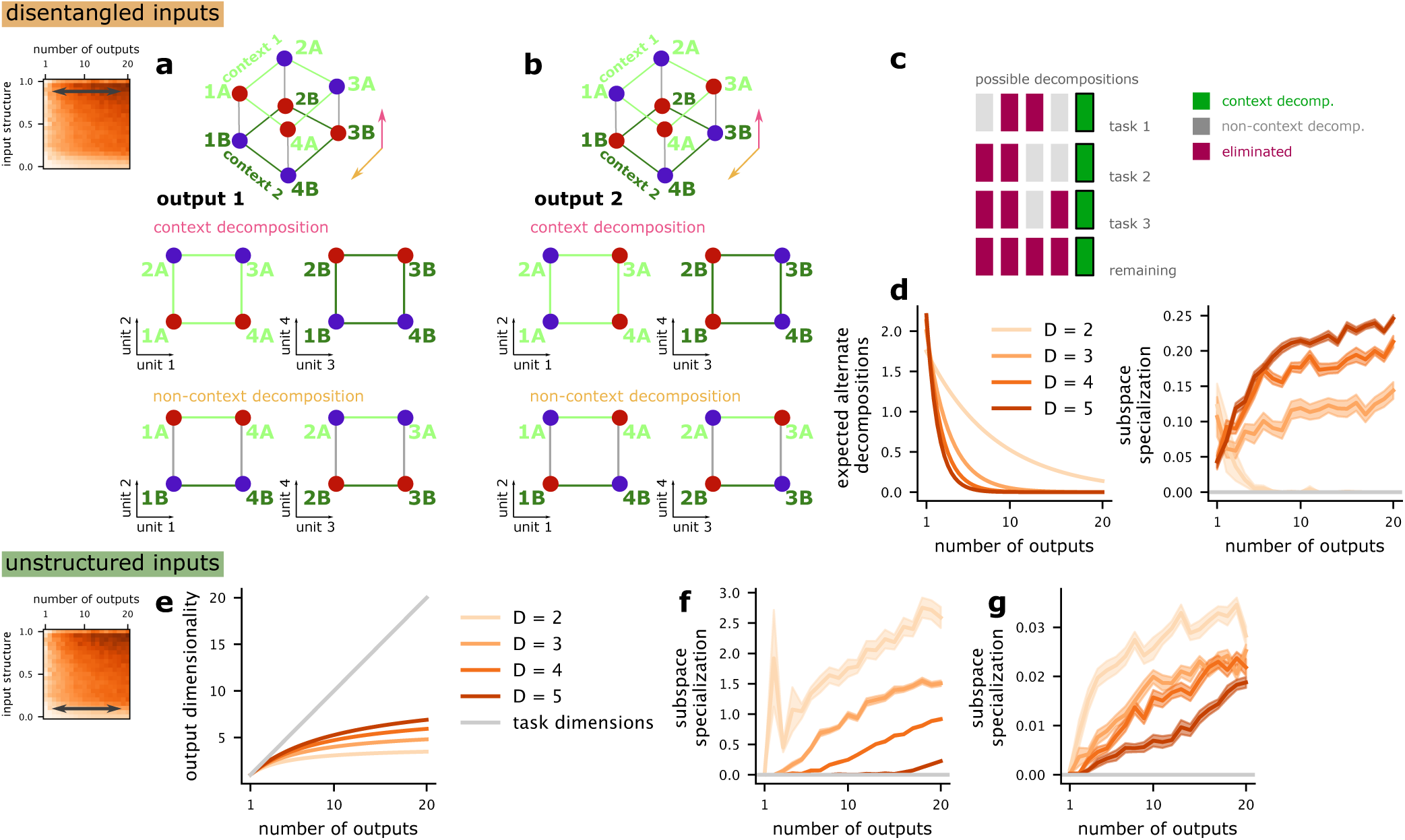
Understanding the emergence of implicit modularity. **a** (top) An example contextual task in a *D* = 2-dimensional latent variable space with a single context variable. (middle) The context decomposition for this task, which makes both component tasks linearly separable. (bottom) An example non-context decomposition which makes the component tasks linearly separable. **b** The same as **a** except the non-context decomposition does not yield linearly separable tasks. Thus a network learning to perform both this task and the task in **a** would be forced to use the context-decomposition. **c** Schematic showing how each task eliminates different non-context decompositions, while the context decomposition is guaranteed to not be eliminated. **d** (left) The expected number of surviving non-context decompositions as a function of the number of output dimensions. (right) The emergence of implicit modularity (i.e., subspace specialization) as a function of the number of output dimensions for different numbers of decision-related variables. The *D* = 2 case goes to zero because implicit modularity is not distinct from the unstructured representation (i.e., both are tetrahedrons). **e** The dimensionality of the required output as a function of the number of output dimensions for different numbers of latent variables. **f** The subspace specialization of the output itself as a function of the number of output dimensions. **g** The subspace specialization in trained complex task networks as a function of number of output dimensions for different numbers of latent variables. The pattern of emergence is the same as in **f**.

Our hypothesis predicts that, once the network begins to learn more quickly (lower average loss) from the unstructured component relative to the disentangled component (fig. S3d, top, grey bar), the modular structure in the learned representation will start to disappear since the unstructured component leads to unstructured representations. Our analysis shows that the points at which Δ learning crosses zero (vertical lines) correspond to the start of a decrease in explicit modularity and a transition to an unstructured or implicitly modular solution (fig. S3d, bottom, grey bar).

We further hypothesize that the transition point at which the unstructured component begins to provide faster learning will depend on the number of irrelevant latent variables. In particular, as the number of irrelevant latent variables increases, we expect that learning from the unstructured component will slow – since the learning problem itself becomes higher-dimensional and the error gradient used in learning is distributed across more dimensions. In contrast, the disentangled component of the representation will be unaffected or weakly affected, since the relevant part of the representation has the same dimensionality regardless of the number of irrelevant variables. Simulations with different numbers of irrelevant latent variables follow the expected pattern (fig. S3e, different green lines). The amount of modularity begins decreasing for higher input structure in situations with fewer irrelevant latent variables, indicating that this increased learning speed in the unstructured component is affecting the learned representation.

Finally, we visualize the structure of learned representations for three key points along the spectrum of input geometries (fig. S3d, circles along the curve) and two different numbers of irrelevant variables (fig. S3e, light and dark green lines). First, we show that modular structure emerges with high input structure, when the disentangled component dominates learning (fig. S3f, left). Second, we show that this modular structure becomes less apparent as the input structure decreases and the unstructured component begins to dominate learning (fig. S3f, center). Third, in a highly unstructured case, the modular structure is completely eliminated (fig. S3f, right). These results provide an explanation for when explicit modularity emerges as input structure becomes stronger, and indicate that other elements of the input – i.e., the number of irrelevant variables – can affect this transition.

### S4 The model develops an abstract and high-dimensional representational geometry

Next, we study the geometry of representations developed by the complex task network across the entire hidden layer for stimuli that are sampled from a single context. We begin by visualizing the activity for the same range of parameters discussed above (fig. S4). We show that increasing the number of output dimensions for the contextual tasks also increases the disentangled structure of the representations, consistent with previous work[3]. In particular, when the network is trained to perform a single task, the dimension relevant to that task dominates the within-context representation (fig. S4a, left). These task-specific representations emerge when animals are overtrained on specific categorization tasks as well[4–6]. In contrast, when the network is trained to use multiple output dimensions within each context, the representation preserves information about all of the relevant latent variables within each context (fig. S4a, right), consistent with previous work in artificial neural networks[3]. These variables are encoded in low dimensional disentangled representation, which reconstructs the part of the latent space containing the relevant variables.

To quantify this change in structure, we adapt the cross-condition generalization performance used in previous work[3, 7] to quantify disentangled or abstract structure in neural representations. Within each context, we train a linear decoder to perform a random, linear classification that depends on the relevant latent variables. However, we train this decoder only using a subset of all stimulus conditions (fig. S4b, “train”), and then test whether or not it generalizes to the held-out set of stimulus conditions (fig. S4b, “test”; see *Cross-condition generalization performance* in *Methods* for more details). We then quantify the average generalization performance in the same parameter range as before (fig. S4c) as well as compute the average *D^′^* of the margin of the classifier (fig. S4d). This reveals that generalization performance primarily depends on the number of output dimensions for the contextual tasks and not on the geometry of the input – which is consistent with previous work in artificial neural networks for non-contextual tasks[3]. The average margin *D^′^*, however, strongly reflects the input geometry (fig. S4d), with higher *D^′^* for more structured input representations – which indicates a greater robustness to noise. The high generalization performance even for relatively unstructured learned representations (fig. S4c, d, bottom left of parameter space) indicates that task training still influences representations enough to impose abstract structure prior to the emergence of context specialization.

**Figure S3:**
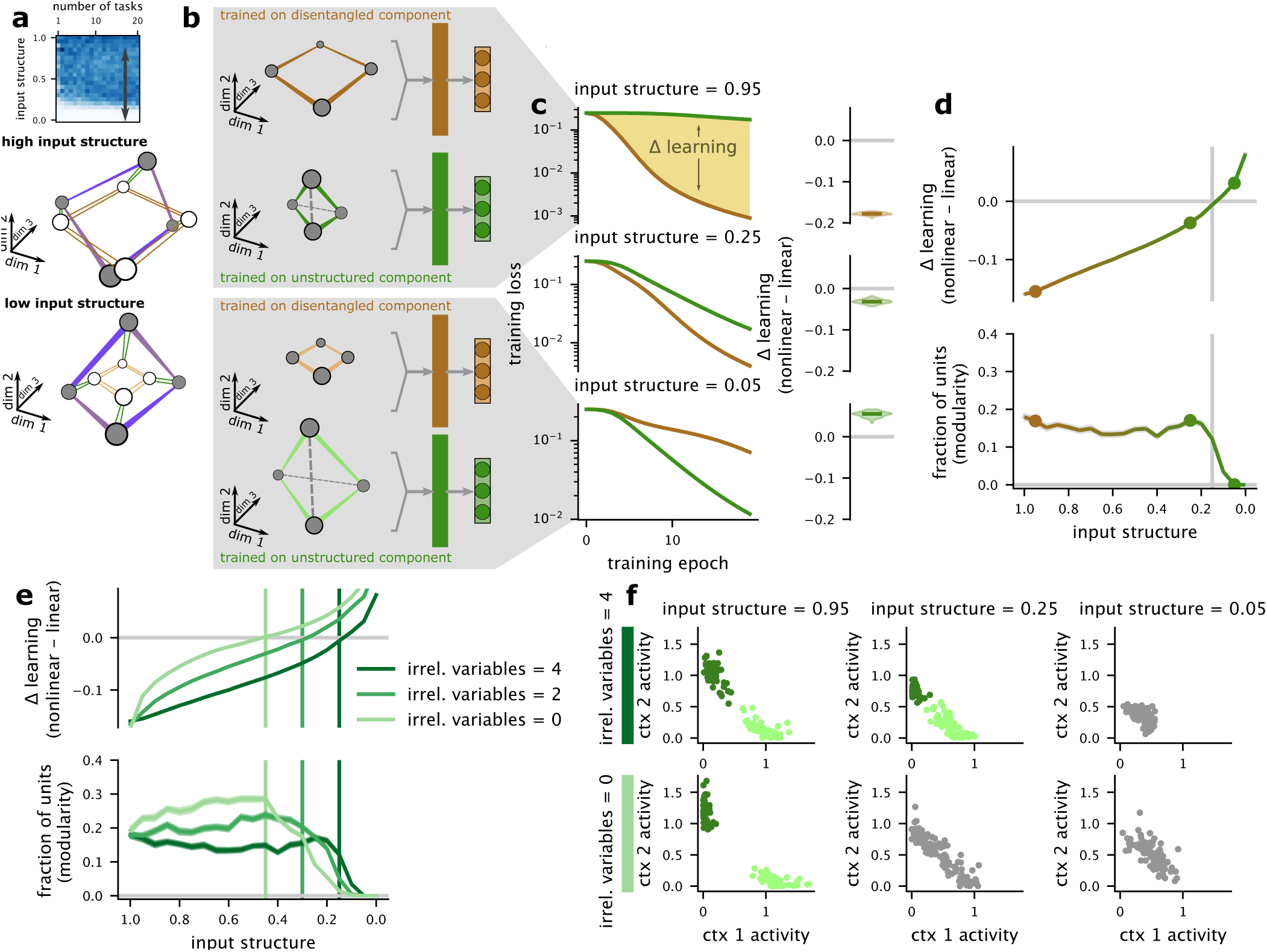
Understanding the emergence of explicit modularity. **a** (top) We focus on the how the representation changes with changes in input structure and contrast between input geometries with high (middle) and low (bottom) structure. **b** All input geometries can be decomposed into distinct disentangled and unstructured components. We decompose input geometries into these two components and train complex task networks on only the isolated component (right). **c** We track the loss of models trained only on the disentangled and unstructured components across training (left), and summarize the average difference (right) as a function of the input structure (top to bottom). **d** As the input structure decreases, the unstructured component begins to yield faster learning than the disentangled component (top). The modular structure learned by the full network starts to disappear (bottom) as the unstructured component begins to provide faster learning (vertical line). **e** The same as **d** for networks with different numbers of irrelevant variables. Fewer irrelevant variables leads to an earlier transition away from modularity. **f** Example average activity in each context for networks with input structure indicated by the outlined points in **d** and either 4 irrelevant variables (top) or 0 irrelevant variables (bottom). While distinct clusters exist for high input structure (left), the transition to faster learning from the unstructured component of the representation is associated with a loss of distinct clusters (center) and eventually with a fully unstructured representation (right).

Further, we quantify the strength of nonlinear perturbations to these low-dimensional abstract representations that emerge as the number of output dimensions for the contextual tasks are increased. To quantify this structure, we use a previously developed measure of representational geometry referred to as the shattering dimensionality (fig. S4e, and see *Shattering dimensionality* in *Methods* for details)[7–9]. A high shattering dimensionality (that is, close to 1) indicates that the representation has the maximum dimensionality for the number of conditions. A high shattering dimensionality requires nonlinear interactions between the representations of different decision-related variables[9]. We find a high shattering dimensionality across the entire range of parameters explored here (fig. S4f) as well as relatively large average margin *D^′^* (fig. S4g). This is surprising because the network does not require any nonlinear interactions between variables to perform tasks within a particular context. Instead, the network must either inherit these nonlinear interactions from the input geometry, or – in the case where the input geometry has no nonlinear interactions – they must emerge when the input representation is through the initially random weights leading to the nonlinear units in the hidden layer.

**Figure S4:**
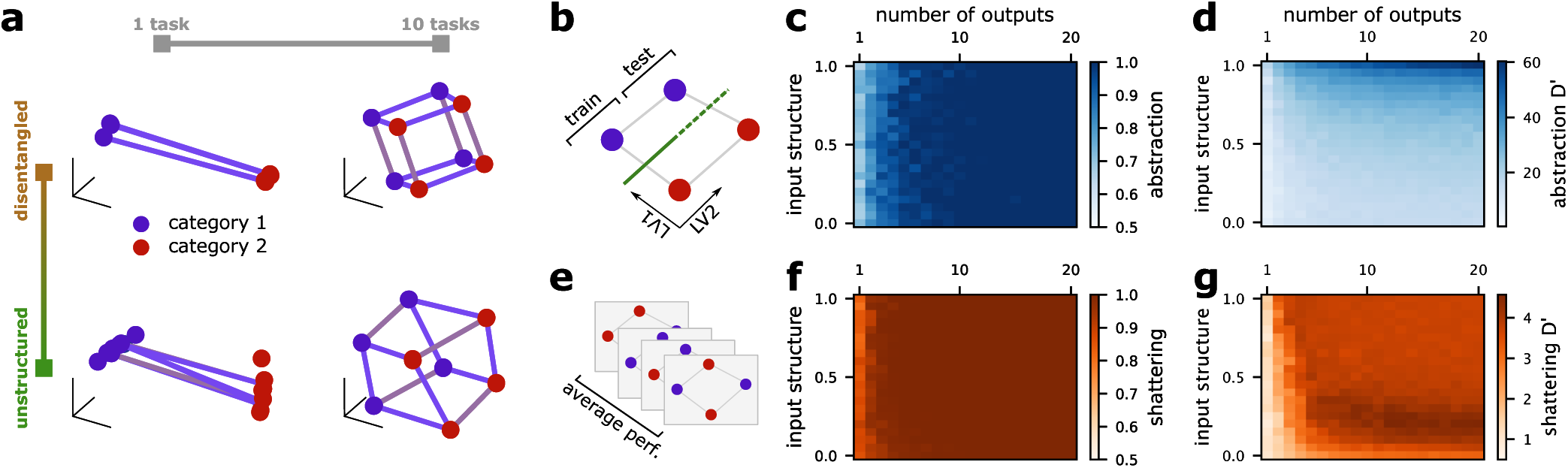
The trained networks have representational geometries within each context that depend on their training. **a** A grid of example models. The plot shows the geometry of representations within each context. The models trained to perform only one task develop a highly specialized geometry for that task. The models trained to perform contextual tasks with more output dimensions represent the full set of latent variables relevant in that context. Each plot shows the representation within a single context, for which three latent variables are relevant. Eight points are shown in each plot. **b** Quantification of the level of abstraction in the representation within each context. Abstraction is measured with the cross-condition generalization performance, where a linear classifier is trained to perform a linear classification when one irrelevant feature is fixed at one value (“trained”) and then tested when the irrelevant feature has the opposite value (“tested”). **c** The level of abstraction of models trained on different numbers of output dimensions and for different input geometries. For a sufficiently large number of output dimensions, the representations are always abstract, no matter the level of entanglement of the inputs. **d** The same as **c** but showing the average *D^′^* of the margin of the classifier for the generalization analysis. The average *D^′^* reflects both the input geometry and number of output dimensions. **e** The shattering dimensionality of representations in each context, measured by quantifying the performance of a linear decoder trained on random, often nonlinear categorizations of all the stimuli in the context. **f** The average performance of a classifier trained on the random classification tasks shown in **e** (i.e., the shattering dimensionality) for models with different input geometries and that were trained with different numbers output dimensions. The shattering dimensionality is always above chance. **g** The same as **f** but showing the average margin *D^′^* of the classifier.

Finally, we investigate how the zero-shot generalization performance of our networks depends on both the input geometry and the number of output dimensions. We train the network to perform contextual tasks as before while holding the value of one irrelevant variable constant (fig. S5a, left). Then, we test the performance of the network when the value of that variable is changed (fig. S5a, right). First, we visualize the representational geometry of the learned configuration in a three-dimensional subspace found by PCA applied only to the trained points (fig. S5b, “trained”). Then, we project the never-before-seen points into the same subspace (fig. S5b, “tested”). We see that unstructured input geometries lead to a far greater change in the representation across learned and novel conditions (fig. S5b, left: low-dimensional; right: high-dimensional). The zero-shot generalization performance of the network is consistent with this visualization. We show that zero-shot performance depends primarily on the geometry of the input representations, rather than the number of output dimensions that the network is trained to perform (fig. S5c). This is because the nonlinear mixing in the input causes the decision variables to be represented in different subspaces given different values for the irrelevant variables.

**Figure S5:**
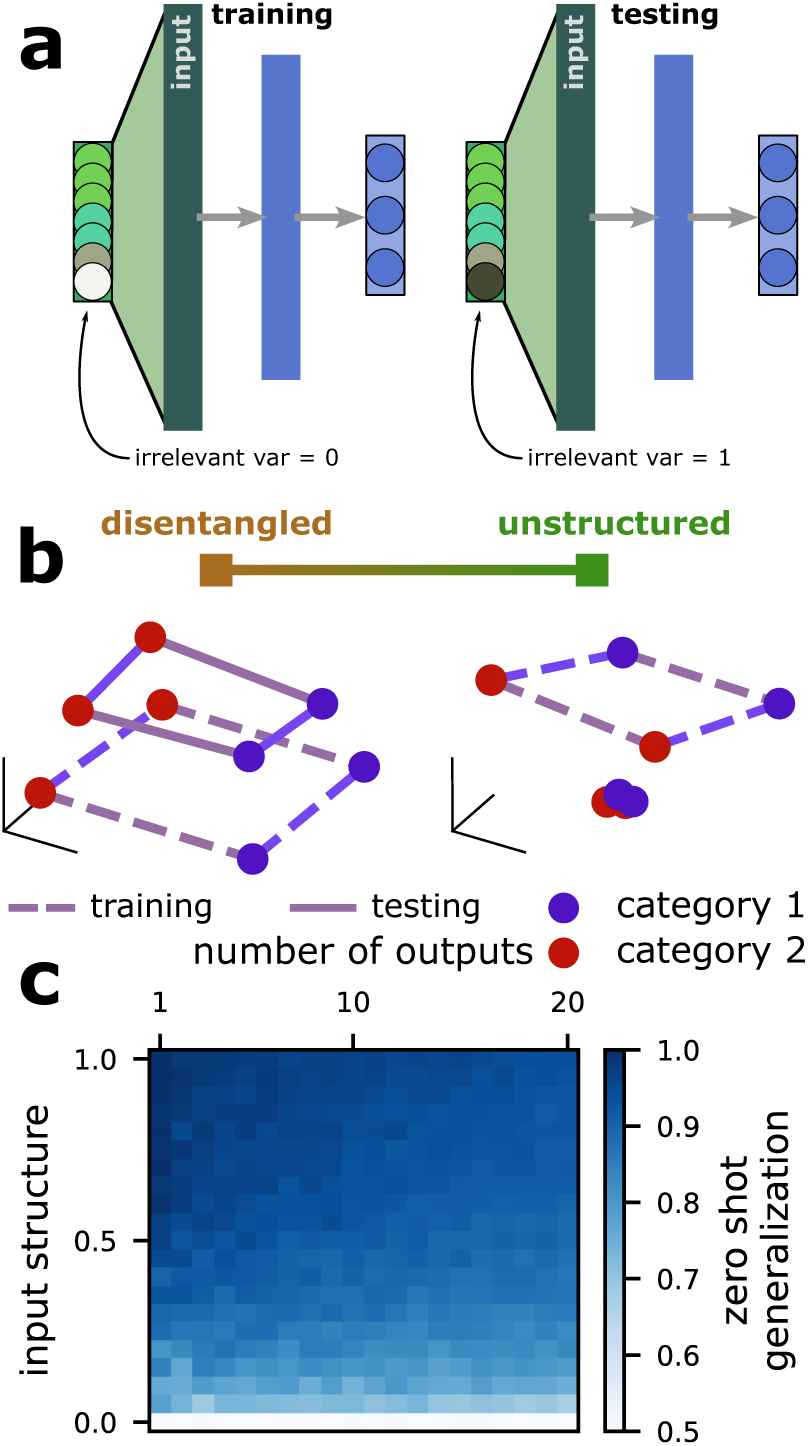
The zero-shot generalization performance of a model depends on the input geometry. **a** Schematic of a zero-shot generalization analysis for the full model, where the full contextual multi-tasking model is trained for a fixed value of an irrelevant variable, then its generalization performance is tested for the opposite value of that variable. **b** Visualization of how representations change within a single context from the learned set of stimuli to the novel set of stimuli, shown for fully disentangled (left) and unstructured (right) input geometries. **c** Zero-shot generalization performance of the models trained and tested with the procedure in **a**, shown for models trained on different numbers of output dimensions and with different input geometries.

### S5 Random coloring tasks

We introduce an additional task variation, random coloring tasks (fig. S6a). In these tasks, each stimulus condition is randomly labeled while ensuring balance between the two categories (fig. S6b, left). In a second variation, we train networks to perform random coloring tasks on distinct subsets of latent variables in different contexts (fig. S6b, right). Here, we find that selectivity in the random coloring task without contextual structure is similar to that which emerges in the arbitrary second order tasks from above, with a clear organization emerging for disentangled inputs (fig. S6c, left) but not for unstructured inputs (fig. S6d, left). The contextual coloring task follows our results from contextual tasks before, where clear context specialization emerges for disentangled inputs (fig. S6c, right) and not for unstructured inputs (fig. S6c, right).

**Figure S6:**
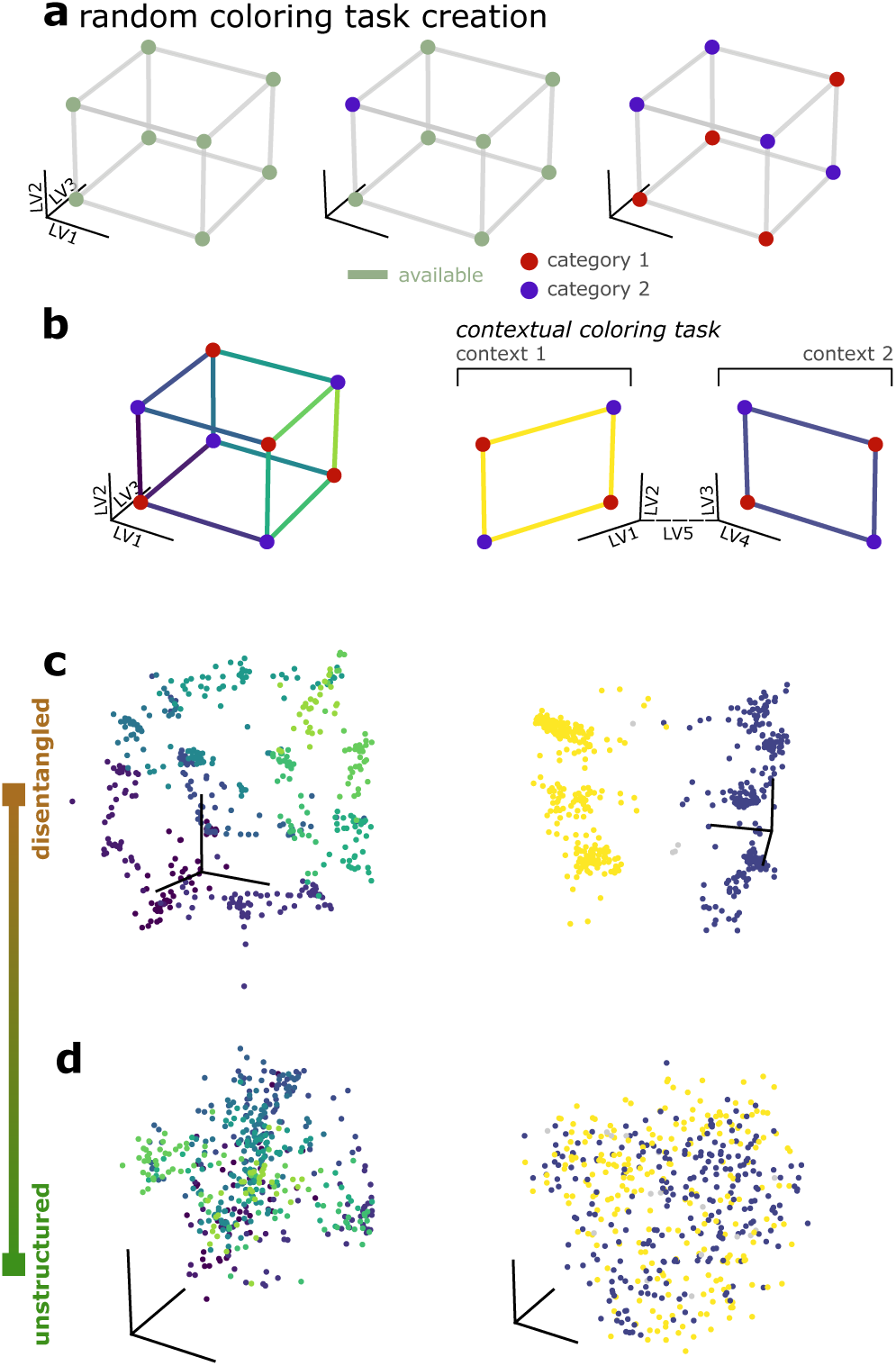
Random, unstructured tasks can also give rise to explicit modularity. **a** The procedure for creating a random coloring task. **b** An example of a random coloring task on a cube (left) and a contextual random coloring task (right). **c** The weight vectors from a network with disentangled input geometry that has been trained to perform a random coloring task (left) and a contextual random coloring task (right) with 200 outputs projected into the input space. **d** The same as **c** except for a network with fully unstructured input geometry.

## References

1. Hirokawa, J., Vaughan, A., Masset, P., Ott, T. & Kepecs, A. Frontal cortex neuron types categorically encode single decision variables. Nature 576, 446–451 (2019).

2. Hocker, D. L., Brody, C. D., Savin, C. & Constantinople, C. M. Subpopulations of neurons in lOFC encode previous and current rewards at time of choice. Elife 10, e70129 (2021).

3. Lee, J. J., Krumin, M., Harris, K. D. & Carandini, M. Task specificity in mouse parietal cortex. Neuron 110, 2961–2969 (2022).

4. Sun, W. et al. Learning produces a hippocampal cognitive map in the form of an orthogonalized state machine. bioRxiv, 2023–08 (2023).

5. Hardcastle, K. et al. A Multiplexed , Heterogeneous , and Adaptive Code for Navigation in Medial Entorhinal Cortex. Neuron 94, 1–13. issn: 0896-6273. 10.1016/j.neuron.2017.03.025 (2017).

6. Posani, L., Wang, S., Muscinelli, S. P., Paninski, L. & Fusi, S. Rarely categorical, always high-dimensional: how the neural code changes along the cortical hierarchy. bioRxiv, 2024–11 (2025).

7. O’Neill, P.-K. et al. The representational geometry of emotional states in basolateral amygdala. bioRxiv, 2023–09 (2023).

8. Boyle, L. M., Posani, L., Irfan, S., Siegelbaum, S. A. & Fusi, S. Tuned geometries of hip-pocampal representations meet the computational demands of social memory. Neuron 112, 1358–1371 (2024).

9. Nogueira, R., Rodgers, C. C., Bruno, R. M. & Fusi, S. The geometry of cortical representations of touch in rodents. Nature Neuroscience 26, 239–250 (2023).

10. Bernardi, S. et al. The geometry of abstraction in the hippocampus and prefrontal cortex. Cell 183, 954–967 (2020).

11. Courellis, H. S. et al. Abstract representations emerge in human hippocampal neurons during inference. Nature, 1–9 (2024).

12. Johnston, W. J., Palmer, S. E. & Freedman, D. J. Nonlinear mixed selectivity supports reliable neural computation. PLoS computational biology 16, e1007544 (2020).

13. Johnston, W. J., Fine, J. M., Yoo, S. B. M., Ebitz, R. B. & Hayden, B. Y. Semi-orthogonal subspaces for value mediate a binding and generalization trade-off. Nature Neuroscience, 1–13 (2024).

14. Konkle, T. Emergent organization of multiple visuotopic maps without a feature hierarchy. bioRxiv, 2021–01 (2021).

15. Bullmore, E. & Sporns, O. The economy of brain network organization. Nature reviews neuroscience 13, 336–349 (2012).

16. Bakhtiari, S., Mineault, P., Lillicrap, T., Pack, C. & Richards, B. The functional specialization of visual cortex emerges from training parallel pathways with self-supervised predictive learning. Advances in Neural Information Processing Systems 34, 25164–25178 (2021).

17. Strotzer, M. One century of brain mapping using Brodmann areas. Clinical Neuroradiology 19, 179 (2009).

18. Van Essen, D. C. & Maunsell, J. H. Hierarchical organization and functional streams in the visual cortex. Trends in neurosciences 6, 370–375 (1983).

19. Rakic, P. Specification of cerebral cortical areas. Science 241, 170–176 (1988).

20. Khosla, M., Williams, A. H., McDermott, J. & Kanwisher, N. Privileged representational axes in biological and artificial neural networks. bioRxiv, 2024–06 (2024).

21. Rigotti, M. et al. The importance of mixed selectivity in complex cognitive tasks. Nature 497, 585–590 (2013).

22. Mante, V., Sussillo, D., Shenoy, K. V. & Newsome, W. T. Context-dependent computation by recurrent dynamics in prefrontal cortex. nature 503, 78–84 (2013).

23. Siegel, M., Buschman, T. J. & Miller, E. K. Cortical information flow during flexible sensori-motor decisions. Science 348, 1352–55. issn: 0036-8075. arXiv: arXiv:1011.1669v3. http://www.sciencemag.org/content/348/6241/1352.full (2015).

24. Driscoll, L., Shenoy, K. & Sussillo, D. Flexible multitask computation in recurrent networks utilizes shared dynamical motifs. bioRxiv (2022).

25. Yang, G. R., Joglekar, M. R., Song, H. F., Newsome, W. T. & Wang, X.-J. Task representations in neural networks trained to perform many cognitive tasks. Nature neuroscience 22, 297–306 (2019).

26. Flesch, T., Juechems, K., Dumbalska, T., Saxe, A. & Summerfield, C. Orthogonal representations for robust context-dependent task performance in brains and neural networks. Neuron 110, 1258–1270 (2022).

27. Minxha, J., Adolphs, R., Fusi, S., Mamelak, A. N. & Rutishauser, U. Flexible recruitment of memory-based choice representations by the human medial frontal cortex. Science 368, eaba3313 (2020).

28. Panichello, M. F. & Buschman, T. J. Shared mechanisms underlie the control of working memory and attention. Nature 592, 601–605 (2021).

29. Xie, Y. et al. Geometry of sequence working memory in macaque prefrontal cortex. Science 375, 632–639 (2022).

30. Dubreuil, A., Valente, A., Beiran, M., Mastrogiuseppe, F. & Ostojic, S. The role of population structure in computations through neural dynamics. Nature neuroscience 25, 783–794 (2022).

31. Fusi, S., Miller, E. K. & Rigotti, M. Why neurons mix: high dimensionality for higher cognition. Current opinion in neurobiology 37, 66–74 (2016).

32. Saxe, A., Sodhani, S. & Lewallen, S. J. *The neural race reduction: Dynamics of abstraction in gated networks* in *International Conference on Machine Learning* (2022), 19287–19309.

33. Chang, L. & Tsao, D. Y. The code for facial identity in the primate brain. Cell 169, 1013–1028 (2017).

34. She, L., Benna, M. K., Shi, Y., Fusi, S. & Tsao, D. Y. Temporal multiplexing of perception and memory codes in IT cortex. Nature, 1–8 (2024).

35. Elsayed, G. F., Lara, A. H., Kaufman, M. T., Churchland, M. M. & Cunningham, J. P. Reorganization between preparatory and movement population responses in motor cortex. Nature communications 7, 13239 (2016).

36. Johnston, W. J. & Fusi, S. Abstract representations emerge naturally in neural networks trained to perform multiple tasks. Nature Communications 14, 1040 (2023).

37. Fascianelli, V. et al. Neural representational geometries reflect behavioral differences in monkeys and recurrent neural networks. Nature Communications 15, 6479 (2024).

38. Higgins, I., et al. Scan: Learning hierarchical compositional visual concepts. arXiv preprint arXiv:1707.03389 (2017).

39. Moulavi, D., Jaskowiak, P. A., Campello, R. J., Zimek, A. & Sander, J. *Density-based clustering validation* in *Proceedings of the 2014 SIAM international conference on data mining* (2014), 839–847.

40. Farrell, M., Recanatesi, S. & Shea-Brown, E. From lazy to rich to exclusive task representations in neural networks and neural codes. Current Opinion in Neurobiology 83, 102780 (2023).

41. Dasgupta, S. & Gupta, A. An elementary proof of a theorem of Johnson and Lindenstrauss. Random Structures & Algorithms 22, 60–65 (2003).

42. Ganguli, S. & Sompolinsky, H. Compressed sensing, sparsity, and dimensionality in neuronal information processing and data analysis. Annual review of neuroscience 35, 485–508 (2012).

43. Giryes, R., Sapiro, G. & Bronstein, A. M. Deep neural networks with random gaussian weights: A universal classification strategy? IEEE Transactions on Signal Processing 64, 3444–3457 (2016).

44. Cho, Y. & Saul, L. K. Large-margin classification in infinite neural networks. Neural computation 22, 2678–2697 (2010).

45. Shan, H., Minni, S. & Duncker, L. Separating the what and how of compositional computation to enable reuse and continual learning. arXiv *preprint arXiv:2510.20709* (2025).

46. Okazawa, G. & Kiani, R. Neural Mechanisms that Make Perceptual Decisions Flexible. Annual Review of Physiology 85, 191–215 (2023).

47. Crapse, T. B., Lau, H. & Basso, M. A. A role for the superior colliculus in decision criteria. Neuron 97, 181–194 (2018).

48. Hanks, T. D., Mazurek, M. E., Kiani, R., Hopp, E. & Shadlen, M. N. Elapsed decision time affects the weighting of prior probability in a perceptual decision task. Journal of Neuroscience 31, 6339–6352 (2011).

49. Tye, K. M. et al. Mixed selectivity: Cellular computations for complexity. Neuron 112, 2289– 2303 (2024).

50. Litwin-Kumar, A., Harris, K. D., Axel, R., Sompolinsky, H. & Abbott, L. Optimal Degrees of Synaptic Connectivity. Neuron 0, 1153–1164.e7. http://linkinghub.elsevier.com/ retrieve/pii/S0896627317300545 (2017).

51. Neath, A. A. & Cavanaugh, J. E. The Bayesian information criterion: background, derivation, and applications. Wiley Interdisciplinary Reviews: Computational Statistics 4, 199–203 (2012).

1. Alleman, M., Lindsey, J. W. & Fusi, S. Task structure and nonlinearity jointly determine learned representational geometry. arXiv preprint arXiv:2401.13558 (2024).

2. Saxe, A., Sodhani, S. & Lewallen, S. J. *The neural race reduction: Dynamics of abstraction in gated networks* in *International Conference on Machine Learning* (2022), 19287–19309.

3. Johnston, W. J. & Fusi, S. Abstract representations emerge naturally in neural networks trained to perform multiple tasks. Nature Communications 14, 1040 (2023).

4. Swaminathan, S. K. & Freedman, D. J. Preferential encoding of visual categories in parietal cortex compared with prefrontal cortex. Nature neuroscience 15, 315–320 (2012).

5. Sarma, A., Masse, N. Y., Wang, X.-J. & Freedman, D. J. Task specific versus generalized mnemonic representations in parietal and prefrontal cortices.

6. Freedman, D. J. & Assad, J. A. Experience-dependent representation of visual categories in parietal cortex. Nature 443, 85 (2006).

7. Bernardi, S. et al. The geometry of abstraction in the hippocampus and prefrontal cortex. Cell 183, 954–967 (2020).

8. Fusi, S., Miller, E. K. & Rigotti, M. Why neurons mix: high dimensionality for higher cognition. Current opinion in neurobiology 37, 66–74 (2016).

9. Rigotti, M. et al. The importance of mixed selectivity in complex cognitive tasks. Nature 497, 585–590 (2013).

